# Towards accurate, reference-free differential expression: A comprehensive evaluation of long-read *de novo* transcriptome assembly

**DOI:** 10.1101/2025.02.02.635999

**Authors:** Feng Yan, Pedro L. Baldoni, James Lancaster, Matthew E. Ritchie, Mathew G. Lewsey, Quentin Gouil, Nadia M. Davidson

## Abstract

Long-read RNA sequencing has significantly advanced transcriptomics by enabling the full length of transcripts to be assessed. However, current analysis methods often depend on a high-quality reference genome and gene annotation. Recently, *de novo* assembly methods have been developed to utilise long-read data in cases where a reference genome is unavailable, such as in non-model organisms. Despite the potential of these tools, there remains a lack of benchmarking and established protocols for optimal reference-free, long-read transcriptome assembly and differential expression analysis.

Here, we comprehensively evaluate the current state-of-the-art long-read *de novo* transcriptome assembly tools, RATTLE, RNA-Bloom2 and isONform, and compare their performance to one of the leading short-read assemblers, Trinity. We assess various metrics, including assembly quality and computational efficiency, across a range of datasets, which include simulated data and spike-in sequin transcripts, where ground truth is known, and real data from human and pea (*Pisum sativum*) samples, using a reference-based approach to define truth. To represent contemporary analysis scenarios, the datasets cover depths from 6 million to 60 million reads, Oxford Nanopore Technologies (ONT) cDNA, ONT direct RNA and Pacific Biosciences (PacBio) 10x single-cell sequencing. Critically, we assessed the downstream impact of assembly choice on the detection of differential gene and transcript expression.

Our results confirm that long reads generate longer assembled transcripts than short-reads for reference-free analysis, though limitations remain compared to reference-guided approaches, and suggest scope for improved accuracy and reduced redundancy. Of the *de novo* pipelines, RNA-Bloom2, coupled with Corset for transcript clustering, was the best performing in terms of both accuracy and computational efficiency. Our findings offer guidance when selecting the most effective strategy for long-read differential expression analysis, when a high-quality reference genome is unavailable.

## Introduction

Understanding which genes and transcripts are present in an organism is essential for analysing RNA sequencing (RNA-seq) data and enables a variety of downstream analyses, such as differential expression, alternative splicing, and the identification of RNA variants. Transcriptome assembly is a key method to achieve this, particularly when reference genomes and gene annotations are unavailable. Transcriptome assembly reconstructs the sequences of expressed transcripts directly from RNA-seq data and is performed using two primary approaches: reference-guided and *de novo*. Reference-guided assembly relies on a reference genome to infer genes and their exon structures based on the positions of mapped reads (Pertea et al., 2015). This method is highly effective, often enabling the identification of novel transcripts even in well-characterised species like humans and mice (Zhao et al., 2023). However, its success relies on the availability and quality of a reference genome. In contrast, *de novo* assembly offers a robust solution for non-model organisms that lack reference genomes. By reconstructing transcriptomes directly from RNA-seq reads, this approach bypasses the need for prior genomic information. It allows gene expression and transcript diversity to be explored in a broad array of species, significantly extending the applicability of RNA-seq.

To achieve *de novo* assembly of massively parallel short-read sequencing, numerous tools and pipelines have been developed such as Trinity, RNA-Bloom, SOAPdenovo-Trans, rnaSPAdes and Oases (Bushmanova et al., 2019; Grabherr et al., 2011; Nip et al., 2020; Schulz et al., 2012; Xie et al., 2014). Collectively they have facilitated the analyses of thousands of transcriptomes (Hölzer & Marz, 2019; Raghavan et al., 2022), expanding our knowledge of the diversity and complexity of life beyond model species. Although generally applied to non-model organisms (Marlétaz et al., 2018; G.-Q. Zhang et al., 2017; Y. Zhang et al., 2024), *de novo* assembly has also proven beneficial to the analysis of cancer transcriptomes, in particular in the detection of gene rearrangements (Cmero et al., 2021; Davidson et al., 2015; Gonzalez-Bosquet et al., 2022; Wang et al., 2022).

However, despite the success of short-read *de novo* transcriptome assembly, a fundamental limitation remains in that long-range relationships across exons and exon junctions can not be directly measured and must be inferred. Due to fragmentation, splicing events at one end of a gene are decoupled from those at the other end, and the true exon structure of the gene is obscured. This limitation is solved by long-read (or third-generation) sequencing. Both Pacific BioSciences (PacBio) and Oxford Nanopore Technologies (ONT) offer protocols to sequence full-length transcripts without fragmentation and ONT is even capable of sequencing native RNA with RNA modifications (Garalde et al., 2018). Both technologies offer the promise of higher-quality assembled transcriptomes. When combined with single-cell technologies, they enable isoform-resolved single-cell transcriptome analysis (Tian et al., 2021).

To leverage these advances, new reference-guided assembly methods have been built for long reads including Bambu, StringTie2, IsoQuant and FLAIR (Chen et al., 2023; Kovaka et al., 2019; Prjibelski et al., 2023; Tang et al., 2020). In contrast, few assembly methods which work without a reference genome or annotation have been developed. The first efforts of such kind were CARNAC-LR and isONclust which cluster reads into genes or gene families (Marchet et al., 2019; Sahlin & Medvedev, 2020). However, these two methods do not generate a set of non-redundant transcript sequences, restricting their application. RATTLE was the first developed to address the transcript assembly problem, which it solved using a three-step pipeline. First, reads are clustered into gene and then transcript groups using a greedy *k*-mer based approach for efficiency. Next, errors are corrected by consensus calling the reads within each transcript cluster. A final polishing step refines the clusters and reports the number of reads in each to give transcript quantification (de la Rubia et al., 2022). RATTLE was shown to improve clustering accuracy and provided comparable transcript abundance estimates to reference-guided tools when tested on SIRV spike-in transcripts. Following RATTLE, RNA-Bloom2 was developed. It builds upon RNA-Bloom, a successful short-read transcriptome assembler, and employs error-tolerant digital normalization to reduce the number of comparisons required for the subsequent steps of overlap assembly and polishing (Nip et al., 2023). RNA-Bloom2 was reported to improve computational efficiency over RATTLE, and provide higher recall when tested on simulation and spike-in data. More recently, isONform was developed. It forms the final step in the isON-pipeline of isONclust, isONcorrect, and isONform, by using minimizer pairs from isONclust gene clusters for graph construction. Bubbles in the graph are iteratively popped to remove errors and the final output is a set of transcript sequences (Petri & Sahlin, 2023). IsONform was reported to have higher recall than RATTLE on simulated and SIRV spike-in data, but was significantly slower to run and its performance was not compared against RNA-Bloom2.

Aside from *de novo* assemblers specifically developed for long-reads, there are also methods that support the hybrid assembly of both long and short read data. These tools generally fall into two main categories: (1) those that use short reads to error-correct long reads before assembling, such as RNA-Bloom2 hybrid mode, and (2) those that use short reads to build an assembly graph and then incorporate long reads for scaffolding gaps and resolving complex structures, such as rnaSPAdes and IDP-denovo (Fu et al., 2018; Prjibelski et al., 2020).

Independent benchmarking efforts are critical to the advancement of transcriptome assembly-based protocols. For short-read data, they have exposed the strengths and weaknesses of different short-read approaches (Hölzer & Marz, 2019), and informed data analysis workflows (Raghavan et al., 2022). Similar efforts are now underway for long-read assembly tools, but these have largely focused on reference-guided approaches. Dong *et al.,* benchmarked six tools using ONT in-silico mixtures of two cells lines with synthetic RNA spike-ins (sequins), and found Bambu had the highest precision and recall of known sequin transcripts and performed well on low sequencing depths (Dong et al., 2023; Hardwick et al., 2016). Similar results were observed by Su *et al*., who benchmarked 9 tools using simulated PacBio and ONT data and sequins from ONT and found that Bambu, IsoQuant and StingTie2 overall performed best in isoform detection (Su et al., 2024). The Long-read RNA-Seq Genome Annotation Assessment Project (LRGASP) presented a comprehensive assessment of 14 tools across three species with various library and sequencing protocols (both long-read and short-read) on three key tasks: isoform detection with high-quality reference genome and gene annotation, isoform quantification, and isoform detection given a genome but no gene annotation (Pardo-Palacios, Wang, et al., 2024). However, only one long-read *de novo* assembler (RNA-Bloom2) was included. It demonstrated high recall but low precision at recovering SIRV spike-in transcripts. Only one independent study has evaluated *de novo* methods thoroughly (Sagniez et al., 2024). Sagniez *et al*., compared RATTLE, RNA-Bloom2 and isONclust assemblers using deeply sequenced synthetic RNA (SIRV and sequins) with ONT R9 and R10 chemistries. As expected, they found that transcript assembly was most accurate when both a reference genome and gene annotation were used, but *de novo* methods did outperform some reference-guided approaches. However, this study only evaluated the assembly of synthetic transcripts, and used datasets which are a fraction of the depth (O(1 million reads)) and complexity (69 SIRV and 160 sequin transcripts) seen for real transcriptomic projects (O(10 million read), >20k transcripts). Moreover, the impact on downstream differential analysis, being one of the most common applications of RNA-Seq, was not examined.

To overcome the lack of independent benchmarking, we have comprehensively evaluated the performance of current *de novo* assembly methods for long-read sequencing data. As the performance of *de novo* methods varies with transcriptome complexity, sequencing error rate and depth, we tested each method on several datasets: simulated ONT cDNA data, ONT_ PCR-cDNA and ONT_direct RNA data from multiple cancer cell lines with additional sequin spike-ins, PacBio Kinnex single-cell RNA-seq of human peripheral blood mononuclear cells (PBMCs), and in a plant species (PCR cDNA data from *Pisum sativum*). For each, we examined the assembly accuracy and completeness, as well as the software’s computational time and memory. As the end goal of many *de novo* analyses is the identification of differentially expressed genes and transcripts, we assessed the impact that the choice of assembly tool, transcript to gene clustering and quantification makes on the detection of differential expression. We demonstrated that *de novo* methods can recover the majority of differentially expressed genes found with reference-guided approaches and some at the transcript-level. Based on our simulation results, long-read approaches are currently on par with short reads for quantification and differential expression, yet provide more complete transcript sequences. We also assessed hybrid short and long-read assembly pipelines and found that they did not offer an advantage over using a single sequencing protocol. Critically, these results provide the foundational guidelines for the optimal workflow to support downstream long-read differential analysis in species lacking a completed reference genome and gene annotation.

## Results

### Study design

To evaluate the performance of long-read *de novo* assembly tools, we selected three recently developed methods: isONform, RATTLE and RNA-Bloom2. As a gold standard for short-read assembly, we included Trinity, which was chosen based on its strong performance in several independent evaluations (Grabherr et al., 2011; Hölzer & Marz, 2019). In addition, all datasets were assembled and expression quantified with the reference-based tool Bambu (Chen et al., 2023). For datasets without available ground truth, i.e., all except for the simulated and spike-in datasets, Bambu was used to define the truth set. Bambu was chosen due to its high accuracy in independent benchmarking studies and thus represents the best-case scenario for assessing the potential of reference-free methods (Dong et al., 2023; Su et al., 2024).

For downstream differential expression analysis of *de novo* assembled transcriptomes, we employed Corset (Davidson & Oshlack, 2014) to group transcripts into gene clusters, minimap2 (Li, 2018) for aligning reads back to the assembled transcriptome. Salmon (Patro et al., 2017) was used for transcript abundance quantification with bootstrapping (Figure 1) for bulk samples, and Oarfish (Zare Jousheghani et al., 2025) for the single-cell data. Transcript and gene-level differential analysis was then conducted using the Bioconductor package limma (Baldoni et al., 2024, 2025; Law et al., 2014; Ritchie et al., 2015).

**Figure 1:**
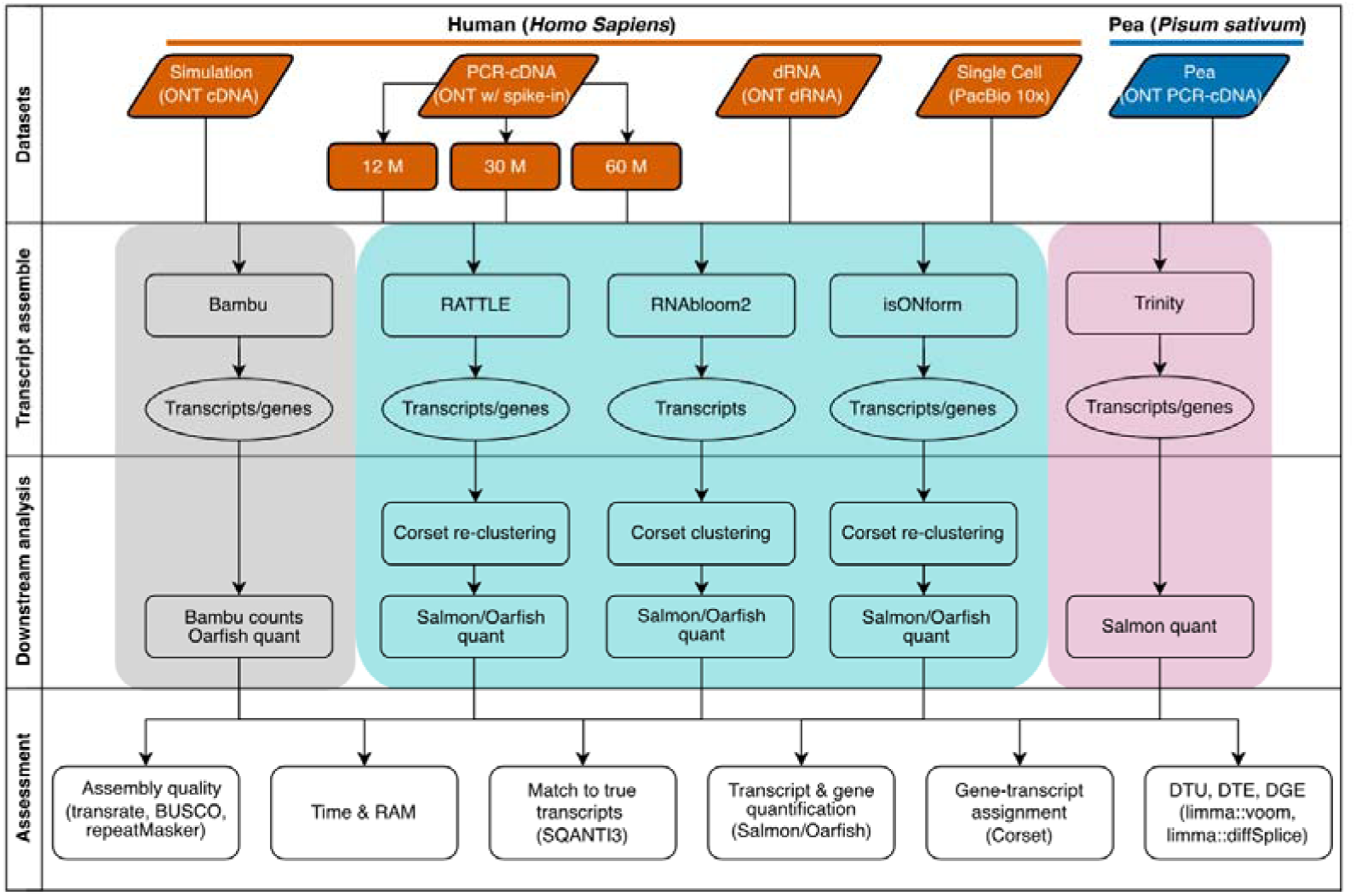
Schematic of the benchmarking study design. Five datasets were utilized, including simulated unstranded cDNA ONT data (simulation), human stranded PCR-cDNA ONT data (PCR-cDNA) with sequin spike-ins, human direct RNA ONT data (dRNA), human single-cell RNA-seq generated with PacBio Kinnex (single-cell), and stranded PCR-cDNA ONT data from Pisum sativum (Pea). The human PCR-cDNA was down-sampled to three sequencing depths (12, 30 and 60 million reads), and included sequin spike-ins (2% of total reads). Each dataset was assembled using three long-read reference free methods: RATTLE, RNA-Bloom2, isONform (teal). Trinity (pink) was applied to matched short-read data when available. A reference-guided approach, Bambu was used to define ground truth (grey). The evaluation focused on both assembly quality and the results of differential gene and transcript expression analysis. Tools used throughout the assembly pipeline and evaluation are also indicated.

We evaluated all methods across five independent datasets, representing various sequencing technologies, read depths, and organisms (Table 1). The datasets included simulated unstranded cDNA ONT data, real PCR-cDNA ONT sequencing data from human lung cancer cell lines with sequin spike-ins (bioinformatically restranded), direct RNA ONT sequencing data from the SG-NEx project, 10x Genomics single-cell RNA-seq generated from the PacBio Kinnex protocol, and PCR-cDNA ONT sequencing data from pea (*Pisum sativum)* cultivars, a non-model species (Chen et al., 2021; Dong et al., 2023; Mestre-Tomás et al., 2023). To investigate the impact of coverage and library size, we downsampled the lung cancer cell line dataset at three different read depths. For all datasets except the single-cell data, we also had matched short-reads available which we downsampled to the same number of nucleotides as the long-read data, and processed with Trinity (Table S1).

**Table 1:** The datasets used for benchmarking. The DE analysis was performed comparing the 2 conditions listed, and the number of true DE genes, DE transcripts and genes undergoing DTU.

| Dataset | Protocol | Total reads (million) | Error rate (%) | Mean read length (bp) (range) | How truth was defined | True #DGE #DTE #DTU-genes | Source |
| --- | --- | --- | --- | --- | --- | --- | --- |
| **Simulation:** control vs perturbed | ONT cDNA (R9.4) | 6 | 7.65 | 1085 (62-44010) | Simulation | 2000, 6933, 2000 | New data generated with SQANTI-SIM |
| **PCR-cDNA:** Human lung cancer cell lines HCC827 vs NCI-H1975 + spike-ins | ONT PCR-cDNA (SQK-PCS109, R9.4) | 12 | 6.81 | 685 (50-71138) | Bambu | 5328, 5362, 635 | (Dong et al., 2023) |
|  |  | 30 |  |  | Bambu | 7506, 8502, 1182 |  |
|  |  | 60 |  |  | Bambu | 9094, 11098, 1726 |  |
|  |  | 2% of total reads |  |  | Sequin spike-in | 44, 95, 28 |  |
| **dRNA:** SG-NEx human cancer cell lines MCF7 vs A549 | ONT direct RNA (SQK-RNA002, R9.4) | 11.6 | 15.9 | 1060 (50-16218) | Bambu | 6280, 7231, 1637 | (Chen et al., 2021) |
| **Single-cell:** Human PBMC myeloid vs lymphoid cells | PacBio Kinnex 10x 3' kit | 15 (assembly) 555 (quantification) | 0.44 | 835 (50-7173) | Bambu | 7352, 23711, 7363 | https://www.pacb.com/connect/datasets/#RNA-datasets |
| **Pea:** (*Pisum sativum*) Thomas Laxton vs Daisy cultivars | ONT PCR-cDNA (SQK-PCB109, R9.4) | 16.5 | 7.75 | 649 (50-11561) | Bambu | 254, 403, 87 | In-house new data |

Aside from the pea cultivars and single-cell data, each dataset consisted of two conditions, each in triplicate, with reads pooled for assembly. For the pea, samples from four conditions were pooled for assembly, but only two conditions contrasted for downstream differential analysis. For the single-cell dataset, we first performed gene expression clustering and compared the two major clusters, which corresponded to myeloid and lymphoid cells. For each dataset, we assessed: (1) computational efficiency, (2) the quality of assembled transcripts, based on transcript length, recall of reference transcripts and BUSCO gene recovery, (3) transcript and gene quantification accuracy, particularly focusing on clustering transcripts into gene clusters, and (4) the precision and recall in detecting differentially expressed genes (DGE), differentially expressed transcripts (DTE), and differential transcript usage (DTU), between the two conditions.

### Computational efficiency

Given the substantial time and memory requirements for *de novo* assembly, we first assessed the computational performance of each tool. All assemblers were run on a high-performance computing cluster where we requested 48 cores and a maximum of 1TB of RAM. RATTLE and isONform demonstrated significantly longer assembly times and higher memory usage compared to RNA-Bloom2 (Figure 2, Table S2). Specifically, RATTLE required between 18 hours and 8 days to finish, utilising 110 to 764 GB of RAM, depending on the dataset. While isONform used less memory (65–190 GB), it was slower, taking 51 hours to 7 days to complete assembly and failed to finish within two weeks on the 60 million reads PCR-cDNA and the pea datasets, thus it was not evaluated on these datasets.

**Figure 2:**
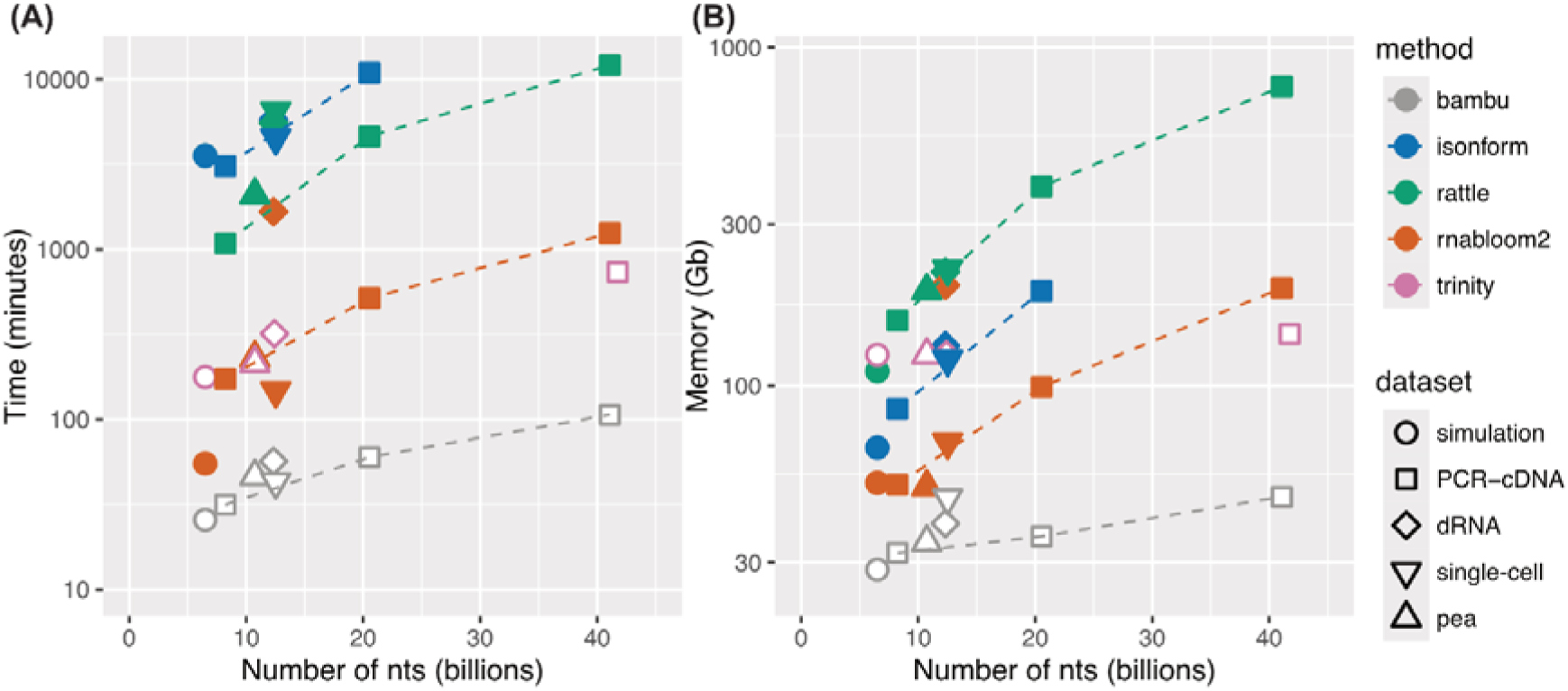
Computational resources required for long read de novo assembly. (A) Time and (B) memory usage of each assembler as a function of the number of total nucleotides for each dataset. The change in resource usage between the three downsampled depths from the PCR-cDNA dataset are indicated with dashed lines. The reference-guided assembler, bambu, is included for comparison and highlights the significant computational resources required for reference-free assembly.

In contrast, RNA-Bloom2 was notably faster and more memory-efficient, completing assemblies in 1 to 27 hours while consuming 50 to 198 GB of RAM. However, its performance on the dRNA dataset was an exception, where it ran 9 times slower and required 4 times more memory compared to cDNA at a similar read depth. We hypothesise that the increased error rate in dRNA sequencing reduced the effectiveness of digital normalisation, which is a key step in RNA-Bloom2 to reduce computational burden (60% of dRNA reads remaining compared to only 12.3% of PCR-cDNA reads after digital normalisation). Trinity, the short-read assembler, which also employs digital normalisation, showed comparable runtime to RNA-Bloom2 across the datasets and scaled consistently with increasing read depth. Notably, Trinity maintained relatively stable memory usage across different read depths.

Overall, we found RNA-Bloom2 to be the most computationally efficient long-read *de novo* assembler, although its performance may degrade on datasets with higher basecalling error rates, such as direct RNA. All *de novo* methods demanded substantial computational resources, highlighting a current limitation in assembling large datasets.

### Assembly quality

*De novo* assemblies from RATTLE, RNA-Bloom2 and isONform, together with Trinity were next assessed for their quality, by comparing characteristics of the assembled transcriptome, such as the number, length and recall of reference transcripts (Figure 3). SQANTI3 was used to link assembled transcripts to reference transcripts using splice junctions (Pardo-Palacios, Arzalluz-Luque, et al., 2024). To assess whether complete transcripts were being assembled, we used the Conditional Reciprocal Best BLAST (CRBB) approach, which gives the percentage of bases recovered for each reference transcript (Aubry et al., 2014; Smith-Unna et al., 2016). As pea is likely to have incomplete gene annotation, we also used BUSCO genes to assess assembly completeness. To set an upper-limit on the best performance expected, we also calculated metrics for the reference genome-guided approach, Bambu, which was filtered for only expressed transcripts (Methods).

**Figure 3:**
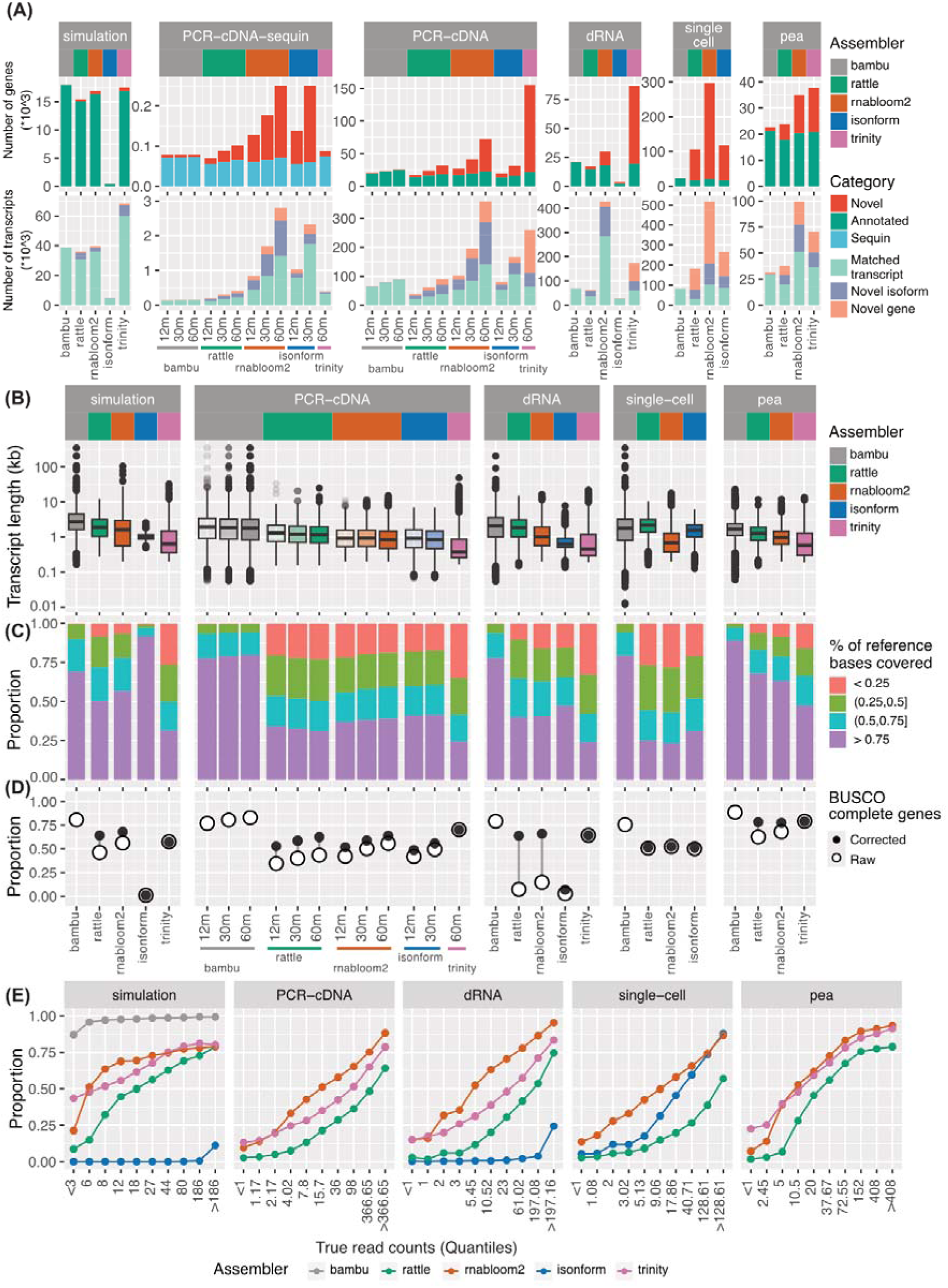
Assessment of assembled transcriptome quality. (A) Number of novel and annotated genes and transcripts in each assembled transcriptome. Transcripts were further divided into matched transcript (complete splice match and incomplete splice match), novel isoform (novel in catalogue and novel not in catalogue) and novel gene (intergenic, genic, intronic, antisense, and fusion). (B) Length distribution of assembled transcriptomes. (C) The proportion of bases recovered for each reference transcript using the Conditional Reciprocal Best BLAST (CRBB) approach. (D) Proportion of BUSCO complete genes (both single copy and duplicated) assembled before and after sequence polishing. Assembled transcripts were corrected using the reference genome to remove errors and assess protein sequence predictions. (E) Proportion of reference transcripts that were assembled binned by true expression. True expression was taken from simulated or Bambu counts, and binned into 10% quantiles. X-axis labels show the corresponding read count range for each quantile. For PCR-cDNA data, only the 60 million data was shown.

The number of transcripts assembled by the *de novo* methods varied widely by assembler and dataset (Figure 3A, S1A, Table S3). In the simulation, RNA-Bloom2 and RATTLE reported a similar number of transcripts (∼40000) and genes (∼17000), which were close to the true number simulated (40509 transcripts and 18145 genes). However, isONform assembled only 4849 transcripts and 444 genes. Combined with similar poor performance on the dRNA dataset, this suggests that isONform with default parameters is not robust to variations in the data, such as error rate (dRNA), strandedness or read length (simulation), which may require adjusting isONform’s parameters such as kmer and window size. Trinity, applied on the matched short-read simulation, assembled a similar number of genes as other tools, but over 60000 transcripts. Although this was significantly more than the number of simulated transcripts, the majority were found to have matching splice junctions with the reference gene annotation. Therefore the larger number of transcripts reported by Trinity is likely a result of a fragmented assembly (64% incomplete splice match), rather than errors. This is further supported by Trinity having the shortest transcript lengths (Figure 3B), and fewest complete transcripts (Figure 3C, S1B) across all datasets, and highlights the benefit of long reads in reconstructing the full length of transcripts.

On the real datasets, we found that the reported number of transcripts was highest from RNA-Bloom2 and lowest in RATTLE, excluding isONform’s failure on dRNA. This high number for RNA-Bloom2 can partially be explained by a high level of transcript redundancy, with multiple assembled contigs matching the same reference transcript (Figure S1C). As expected, we also observed that the number of transcripts increased with read depth in the PCR-cDNA dataset. Although new reference transcripts were assembled at higher depths, so too were novel unannotated sequences. The unannotated novel genes increased 3 and 4 folds in RATTLE and RNA-Bloom2 respectively from 12 million to 60 million reads, and composed the majority of the Trinity assembly. Interestingly, these novel sequences also coincided with a drop in the mean transcript length (Figure 3A-B). We also observed that the assemblies were enriched for simple repeats in the ONT cDNA datasets (PCR-cDNA and pea), and SINE and LINE repeats in the PacBio data (single-cell), hinting at artifacts in either assembly or sequencing protocol, such as representation of intronic regions resulted from internal priming, which are rich in repetitive sequence (Fig S1D, Table S5) (You et al., 2025).

Although we could not definitively know if the novel transcripts were real or false positives, the rates of false positives from the simulation and sequin chromosome (chrIS) in the PCR-cDNA data provided evidence of a high rate of assembly error and redundancy. In the simulation, up to 14% of transcripts and 7% of genes were found to be false positives (Figure 3A, S1A). When focusing on assembled transcripts on chrIS, we observed that for the RNA-Bloom2 assembly on the deepest PCR-cDNA dataset, only 71 out of 251 assembled genes matched to sequin genes, and 1412 out of 2804 assembled transcripts matched to known sequin transcripts (Figure 3A). In addition to false transcripts, we also observed insertion and deletion type errors being incorporated into the sequence of assembled transcripts from ONT data. This impacted the predicted protein translation and hence BUSCO completeness, and was particularly evident in the dRNA data, where up to 50% of conserved BUSCO protein-coding genes expressed were missed (Figure 3D, Table S4). These errors were rescued if a reference genome was provided for polishing the long-read transcriptome assemblies. This issue was not seen for the PacBio or short-read data and suggests that error rate is an important consideration when designing experiments requiring reference-free transcriptome analysis.

As a general trend, RNA-Bloom2 reported more false positives or unannotated transcripts than RATTLE, but also had higher recall of true positives (simulated, spike-in transcripts or reference) (Figure 3A, E), consistent with prior studies (Nip et al., 2023; Pardo-Palacios, Wang, et al., 2024). RATTLE’s more conservative behaviour could be explained by requiring a minimum of 5 reads for consensus calling. However, we found that RATTLE missed transcripts even at higher coverage levels and increasing sequencing depth only improved recovery for lowly expressed genes and transcripts (Figure 3E, S1F). Despite these differences, RNA-Bloom2 and RATTLE had highly similar recall at gene-level, as indicated by the number of recovered reference genes (Figure 3A), transcript base-coverage (Figure 3C), and BUSCO genes (Figure 3D).

### Transcript and gene abundance estimates

Next, we examined how the differences between assemblies impacted transcript and gene-level expression estimates. To compare across different assemblies, we estimated abundances by mapping reads to assembled transcriptomes with minimap2 and applied Salmon or Oarfish for quantification (Methods). For each reference transcript, we identified the best match in the assembly, where the best match was defined as the assembled transcript with the highest expression estimate that was a SQANTI3 splice match. We then assessed the correlation between the estimated and ‘true’ expression, with truth derived from simulated counts, sequin concentration or reference-based quantification.

RNA-Bloom2 transcript counts were consistently one of the most correlated with truth (Figure 4A, S2A, Table S6). IsONform performed the best on the 12 million and 30 million PCR-cDNA data and single-cell data, demonstrating that when it runs successfully, it can exceed the performance of other methods. Trinity performed equally well as RNA-Bloom2 on the simulation, but trailed it on the spike-in dataset. We did not assess Trinity on the real datasets, as the truth was defined from long reads, giving the long-read methods an advantage.

**Figure 4:**
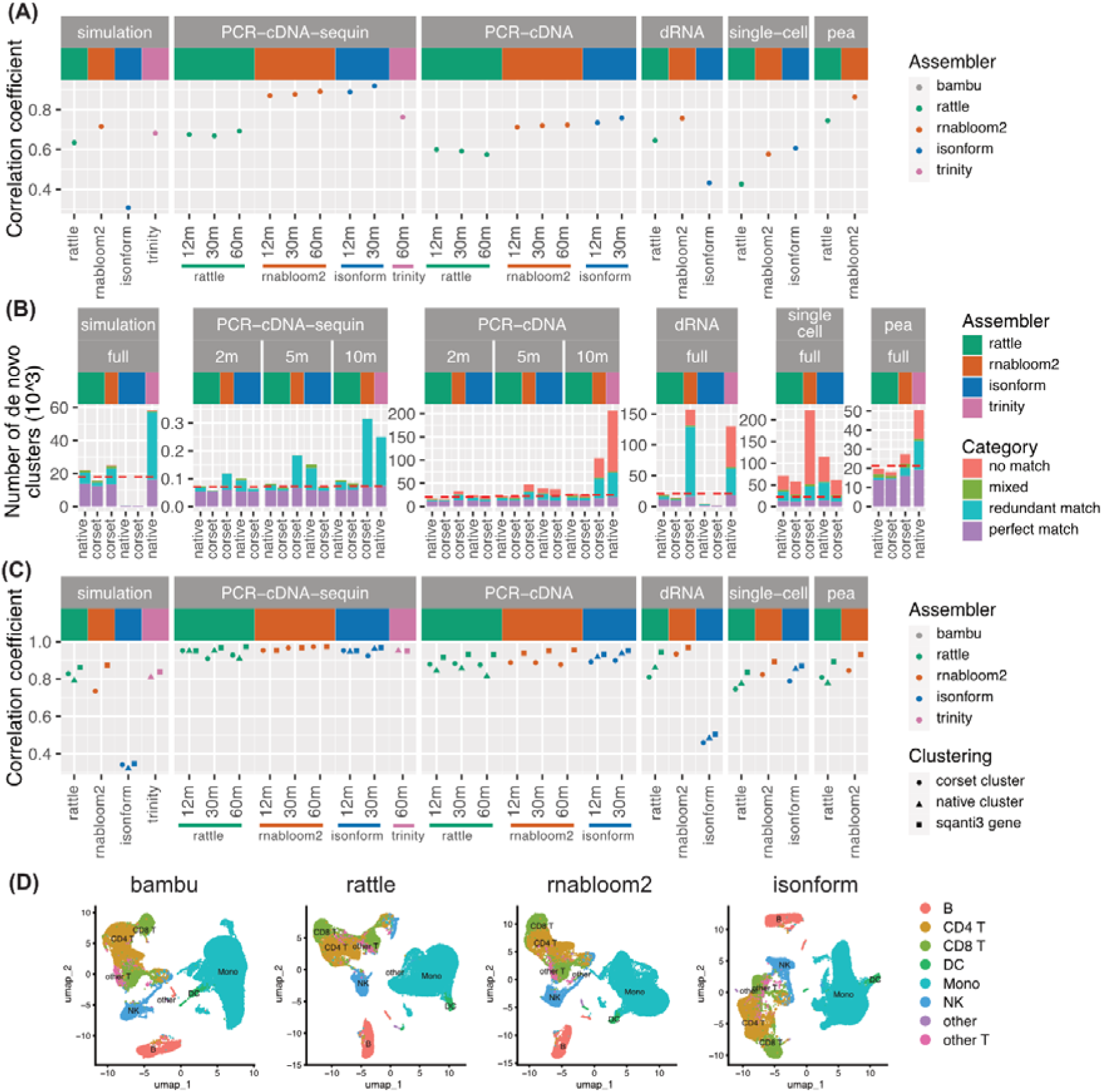
Accuracy of transcript and gene abundance estimates. (A) Pearson correlation coefficients between estimated and true transcripts expression values (log2(count+1)). (B) The number of gene-level clusters generated for each assembly. ‘match’ indicates when all transcripts within the cluster correspond to the same reference gene. Where one reference gene matched multiple clusters, the cluster with the most transcripts was classed as a perfect match and the rest as ‘redundant’. Clusters with transcripts from multiple reference genes were classed as ‘mixed’. Clusters with only novel transcripts were classed as ‘no match’. The red dashed line denotes the number of reference gene-level clusters generated by Bambu. (C) Pearson correlation of true gene expression (log2(count+1)) to Corset, native or SQANTI3 cluster expression. SQANTI3 clustering is an optimal scenario where transcripts are grouped based on their true reference genes, and it is included as an upper limit. (D) UMAP generated using gene-level counts from reference (bambu) and de novo assemblies and colored based on the cell type annotation obtained from a reference analysis.

In order to assess gene-level expression estimates, transcripts must be assigned into gene groups, a step known as clustering. While RATTLE, isONform and Trinity output gene-level clusters, RNA-Bloom2 does not. Therefore, we tested the performance of Corset, which is a clustering method built for short-read *de novo* assembled transcriptomes. Corset clustered the assembled transcripts into genes based on all-vs-all sequence alignments from minimap2. Corset and each assembler’s native clustering were then assessed (Figure 4B, S2B-C, Table S7).

RNA-Bloom2 and Trinity tended to have the highest number of clusters, and a large fraction of these were redundant, meaning there was more than one cluster matching the same reference gene (Figure 4B, S2C, Table S8). For RNA-Bloom2, we found redundant clusters often contained shortened assembled transcripts, with a higher number of sequencing errors, which impacted the all-vs-all alignment (Figure S2D). Both RNA-Bloom2 and Trinity also had many clusters that did not match any reference gene. This is likely a consequence of more novel transcripts being assembled, as seen in the assembly quality assessment (Figure 3A).

We next aggregated the transcript-level counts for each cluster, to obtain gene-level counts, and compared these values against the ‘truth’. Each cluster was assigned a reference gene label based on the reference gene of the majority of its constituent transcripts. When more than one cluster matched a reference gene, we examined the expression of the cluster with the highest counts. When a reference gene had no matching cluster, its count was taken as zero. To assess the impact of clustering, we also looked at the expression correlation for perfect clustering (*SQANTI3 gene*), being the scenario that all assembled transcripts from the same gene are assigned to the same cluster based on SQANTI3 results.

Most *de novo* assemblies gave similar correlations at gene-level (Figure 4C, Table S6). For simulation, RATTLE was slightly better than RNA-Bloom2, while for other datasets, RNA-Bloom2 with Corset had a similar or better performance than RATTLE. Similar to the transcript-level comparison, isONform on the 12 million PCR-cDNA, 30 million PCR-cDNA and single-cell datasets, gave the highest correlation. However, increasing read depth had very little impact on the correlation at both transcript and gene levels.

Overall, Corset was found to be an effective tool for clustering transcripts into genes, with a similar, and in some instances, better performance than native clustering. Based on these results, Corset was used in combination with RNA-Bloom2 to assess downstream analysis steps. The assembler’s own clustering was used for RATTLE and isONform.

Finally, we also examined the impact of abundance estimation on single-cell gene-expression clustering (which is distinct from the transcript clustering examined previously), dimensionality reduction and visualisation. Cell types obtained from a reference analyses were annotated onto the reference-free UMAPs to indicate true groupings. We found that reference-free gene-expression (Figure 4D) and transcript-expression (Figure S3A) were able to recapitulate the clusters seen from Bambu quantification for all assemblers. These results suggest that long-read single-cell analysis can be accurately performed in a reference-free manner, enabling transcriptome analysis at single-cell resolution for non-model species that lack a reference genome.

### Differential expression

We next performed differential analysis using the transcript and gene counts as previously described, and examined each pipeline’s ability to recover true differential gene expression (DGE), differential transcript expression (DTE), and differential transcript usage (DTU) (Baldoni et al., 2024, 2025). True changes were defined using either known simulated or sequin spike-in transcripts and genes when available. For other datasets, we defined truth using a reference-based approach (Bambu) with an FDR cut-off of 0.05 (see Table 1). Transcripts in the *de novo* assembly were assigned to reference transcripts using SQANTI3 and clusters were assigned to reference genes based on majority vote, as described earlier. As benchmarking of differential testing methods has been performed previously (Dong et al., 2023) we did not compare testing methods, but used methods amongst the top performing from prior evaluation, limma::voom and limma::diffSplice (Ritchie et al., 2015) for differential expression and transcript usage analysis respectively.

On the simulated data, all methods performed well in the task of identifying DGE with the true DE genes ranked higher than other clusters (Figure 5A). All methods were close to the accuracy of the reference-guided tool Bambu. Trinity performed marginally worse than the long-read pipelines. When DGE was evaluated using sequin spike-in genes, Trinity and RNA-Bloom2 marginally outperformed others (Figure 5B). On the real datasets, RNA-Bloom2 was consistently a top performer (Figure 5C-F). However when isONForm ran successfully, its performance was similar or exceeded that of RNA-Bloom2. For example, on the PCR-cDNA data, isONform with 30 million reads identified true positives with the same accuracy as RNA-Bloom2 with 60 million reads (Figure 5C). As RNA-Bloom2 produces a highly redundant transcriptome, we also assessed true positives as a function of all top ranked transcripts including duplicated transcripts (Figure S3B-G). Aside from PCR-cDNA DGE, where RATTLE was marginally better, RNABloom2 remained the best performing overall.

**Figure 5:**
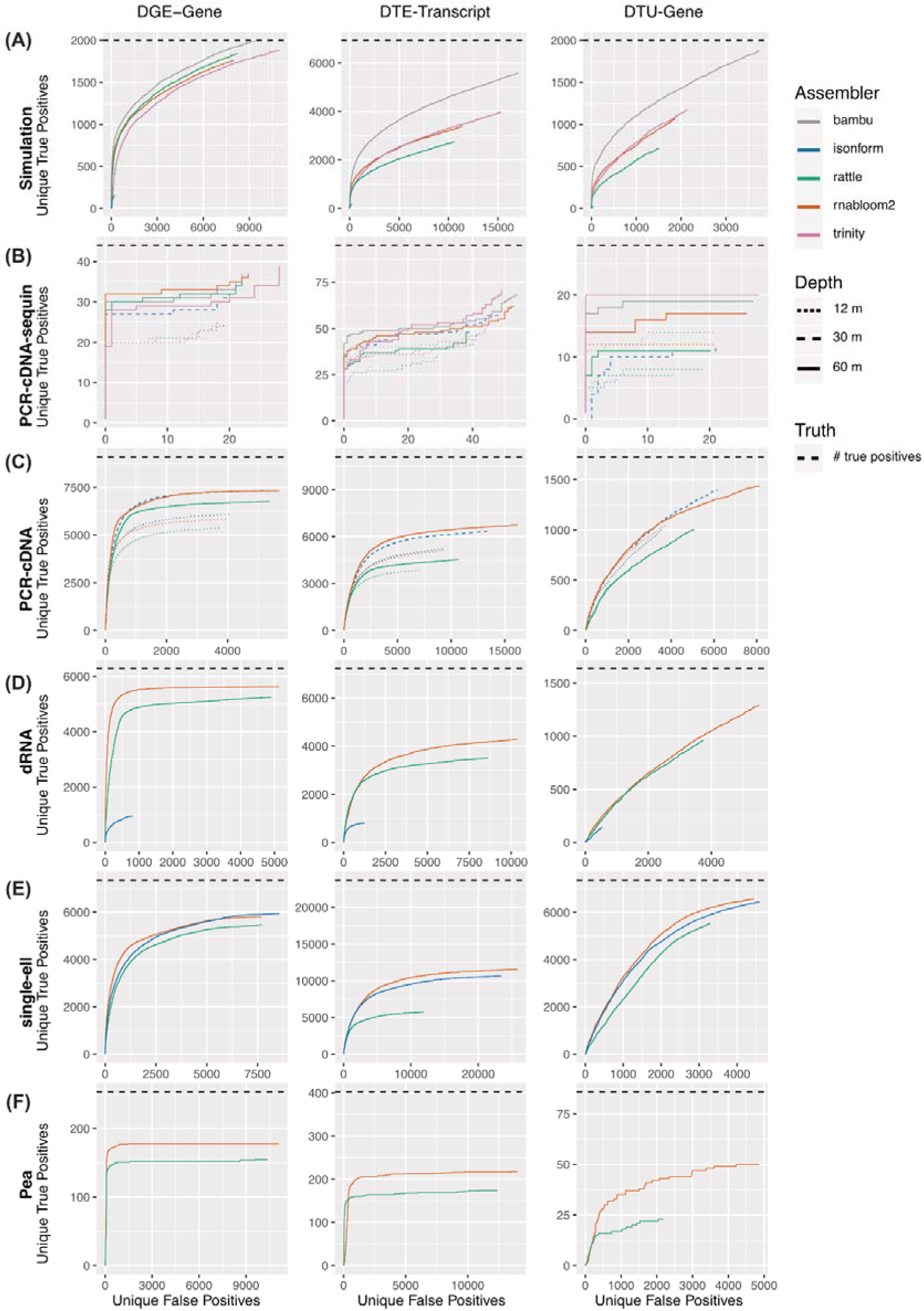
ROC curves for differential analysis. DGE, DTE and DTU analysis was performed for (A) Simulation, (B) PCR-cDNA sequin, (C) PCR-cDNA, (D) dRNA, (E) single-cell, and (F) pea dataset. Unique false positives are shown against unique true positives. When multiple clusters match the same reference gene, the cluster with the lowest FDR is retained, hence ‘unique’ true positives and ‘unique’ false positives. Truth is defined from simulation, sequin or Bambu differential results, and the total number of true positives is indicated by the horizontal dashed line (black). For panels (B-C), we show the deepest and shallowest read depths of the PCR-cDNA dataset for which an assembler ran successfully, specifically 60 million reads for RNA-Bloom2 and RATTLE (solid line), 30 million reads for isONform (lines with longer dashes), and 12 million reads for all assemblers (lines with short dashes).

For DTE, amongst long-read *de novo* assembly methods, RNA-Bloom2 showed the best performance across datasets. Interestingly, the short-read assembler, Trinity performed well on both simulation and sequin data. For differential transcript usage analysis (DTU), genes were ranked by the minimum Simes-adjusted p-values from their transcripts p-values. DTU results showed a similar trend to DGE and DTE, with RNA-Bloom2 being the best performing of the long-read *de novo* assembly methods in 5 out of 6 datasets. We also noticed that for all methods, increasing the read depth in the PCR-cDNA data increased the number of true positives among top ranked results for DGE, DTE and DTU as expected. However, surprisingly, for DTE and DTU, the choice of assembler had a larger impact on performance than quadrupling the sequencing depth (Figure 5B-C).

Interestingly, in real data, all *de novo* methods missed a significant proportion of the differential transcripts reported by Bambu (black dashed line Figure 5C-F). Assuming the Bambu results are true positives, this suggests a significant loss in performance when a reference genome is not used and leaves room for further optimisation of *de novo* methods. However, of the *de novo* assembly tools currently available, RNA-Bloom2 was found to have the best overall performance at both gene and transcript level.

### De novo assembly captures novel transcripts

*De novo* transcriptome assembly enables the discovery of biologically significant transcripts, including those absent from the reference genome and gene annotation, making it an essential approach when the reference genome or gene annotation is incomplete or unavailable. To highlight this advantage, we investigated the differentially expressed de novo assembled transcripts with novel sequences. We considered any transcript to be ‘novel’ if (1) it did not align to reference genome, or (2) its primary alignment to the reference genome contained more than 400 consecutive unaligned bases, or (3) more than 100 or 200 accumulative mismatches, for cDNA and dRNA respectively. Since RNA-Bloom2 performed well on all datasets, we investigated the novel results in its transcriptome assemblies.

In the 60 million PCR-cDNA and dRNA datasets, we identified 2562 and 2307 transcripts respectively which were both differentially expressed and novel compared to the reference transcriptome (Figure 6A-B). A number of these transcripts are likely to contain errors, in particular, some were found to have substantial mismatches likely due to sequencing errors being retained during assembly. However, amongst the top 10 ranked differentially expressed novel transcripts in PCR-cDNA and dRNA, there were three consistent with genomic rearrangement found in the Cancer Cell Line Encyclopedia (CCLE) (Table S9-10) (Ghandi et al., 2019). One of these fuses the RPL7 gene to an intergenic region on chromosome 8 in the HCC827 lung cancer cell line (Figure 6D). The other two were found in the MCF7 breast cancer cell line. One is a fusion of ATXN7 to an intergenic region on chromosome 1 (Figure 6E), and the other is the well-reported fusion BCAS4-BCAS3 (Edgren et al., 2011). To assess whether other fusion genes were seen amongst the novel transcripts, we searched all assembled transcripts for fusions using JAFFAL (Davidson et al., 2022) and compared them to previously reported fusion genes in the CCLE. For PCR-cDNA, 8 novel transcripts were identified as DE fusion transcripts with 4 previously reported fusion genes in CCLE (2 from H1975 and 2 from HCC827, Table S11) (Ghandi et al., 2019). For the dRNA data, 26 novel transcripts were identified as DE fusion transcripts with 11 previously reported fusion genes in CCLE (10 from MCF7 and 1 from the A549 cell line) (Table S12) (Ghandi et al., 2019). When we performed a similar analysis using the novel transcripts from RATTLE, we found 4 DE fusion transcripts in PCR-cDNA, matching 3 CCLE fusion genes, all of which were also detected with RNA-Bloom2. In the dRNA dataset, 9 DE fusion transcripts from RATTLE matched 9 CCLE fusion genes, of which 7 were also detected in RNA-Bloom2 (Figure S4A-B, Table S11-12).

**Figure 6:**
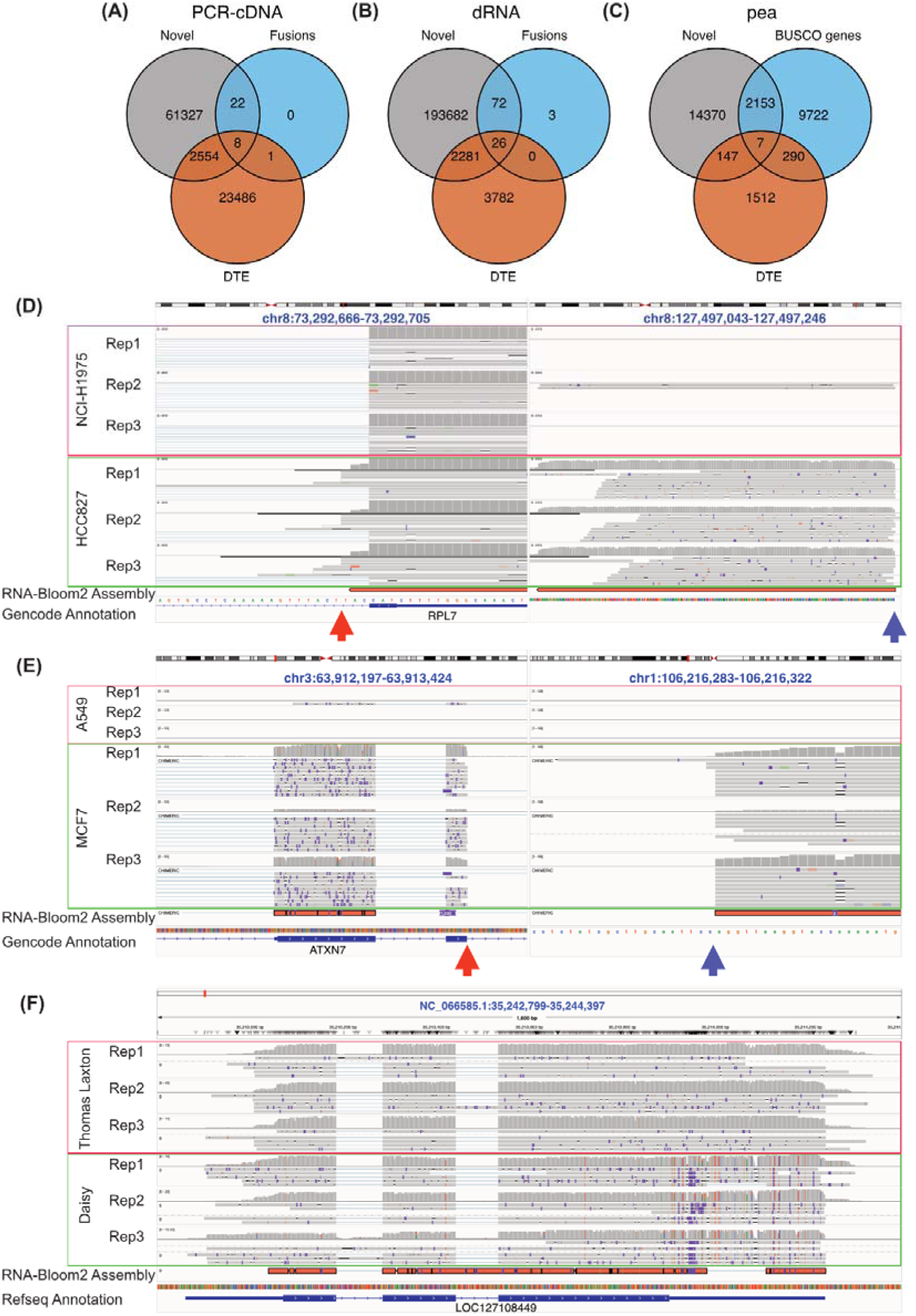
Novel transcripts in de novo assemblies. (A) The number of differentially expressed transcripts by dataset and assembler. Transcripts present in the reference annotation are shown in green, novel but identified via genome-guided assembly in blue, and only in the de novo assembly in red. (B-C) Venn diagram showing the number of transcripts from RNA-Bloom2 which were differentially expressed, novel, and/or known fusion genes in (B) PCR-cDNA 60 million and (C) dRNA. (D) Venn diagram showing the number of transcripts from RNA-Bloom2 which were differentially expressed, novel, and/or has a BUSCO gene match (pea). (E) IGV visualisation of potential RPL7-chr8 intergenic fusion from the PCR-cDNA dataset, split view to show the break point (orange and blue arrows showed left and right breakpoints respectively). (F) IGV visualisation of potential ATXN7-chr1 intergenic fusion from the dRNA dataset, split view to show the break point (orange and blue arrows showed left and right breakpoints respectively). (G) IGV visualisation of LOC127108449 transcript from pea data showed cultivar specific SNPs, insertions and deletions.

For the pea dataset, analysed with RNA-Bloom2, we found 154 DE transcripts with novel sequences, including 7 matching BUSCO genes (Figure 6C, Table S13). These 7 transcripts all showed large sections of divergent sequences when aligned to the reference genome. In addition, these transcripts harboured phased cultivar-specific SNPs. An example of this is in the uncharacterised gene LOC127108449, which has homology with DBH-like monooxygenase in eudicotyledons. We identified a transcript of LOC127108449 which had a small insertion (∼20 bp total), several small deletions and a novel splice variant in its 3’ UTR which was significantly more highly expression in Daisy compared to the Thomas Laxton cultivar (logFC=4.7) (Figure 6F). Read coverage over this loci was consistent with the small novel insertion and deletions in Daisy, but limited reads supported the splice variant, highlighting the need for manual curation of *de novo* assembly results. Interestingly, RATTLE detected more novel DE transcripts matching to BUSCO genes (13 genes), with only 3 matching between the two assemblers (Figure S4C). These results suggest that assembly recall may be improved if multiple assemblies are combined.

These novel examples collectively demonstrate that *de novo* methods do provide insights beyond known genes and transcripts and enable reference-unbiased differential analysis.

### Long-read only pipelines outperform hybrid approaches

Despite the theoretical advantages of long-read sequencing for reference-free differential transcript analysis, our benchmarking results indicate that short reads still perform well. Short-read sequencing currently benefits from lower error rates. It also produces a greater number of counts for a fixed total number of sequenced bases, increasing statistical power for differential testing. Therefore, as a final assessment in our study, we evaluated whether *hybrid* approaches could integrate the strengths of both long- and short-read sequencing to outperform either technology alone.

Using the simulation and 60 million PCR-cDNA datasets, we evaluated two representative hybrid assembly approaches: RNABloom2-hybrid and rnaSPAdes. These two tools represent current mainstream approaches for hybrid assembly. RNABloom2-hybrid extends RNABloom2 by using short-reads to polish the long-read prior to assembly. In contrast, rnaSPAdes uses the short reads to build an assembly graph, then long reads are aligned to the graph to resolve repeats and close gaps.

Each hybrid assembler was applied to a dataset comprising 50% ONT and 50% Illumina reads, with reads randomly selected and the total number of sequenced bases matched to the single-technology datasets. Following assembly, read alignment and quantification were performed separately for long and short reads as described previously, and the resulting count matrices were aggregated to generate the final gene- or transcript-level quantifications. Pipeline performance was then assessed using the same metrics as before, including assembly quality (Figure 7A-C, Figure S5A-B), clustering (Figure 7D-E, S5C-E), and differential expression accuracy (Figure 7F-H).

**Figure 7:**
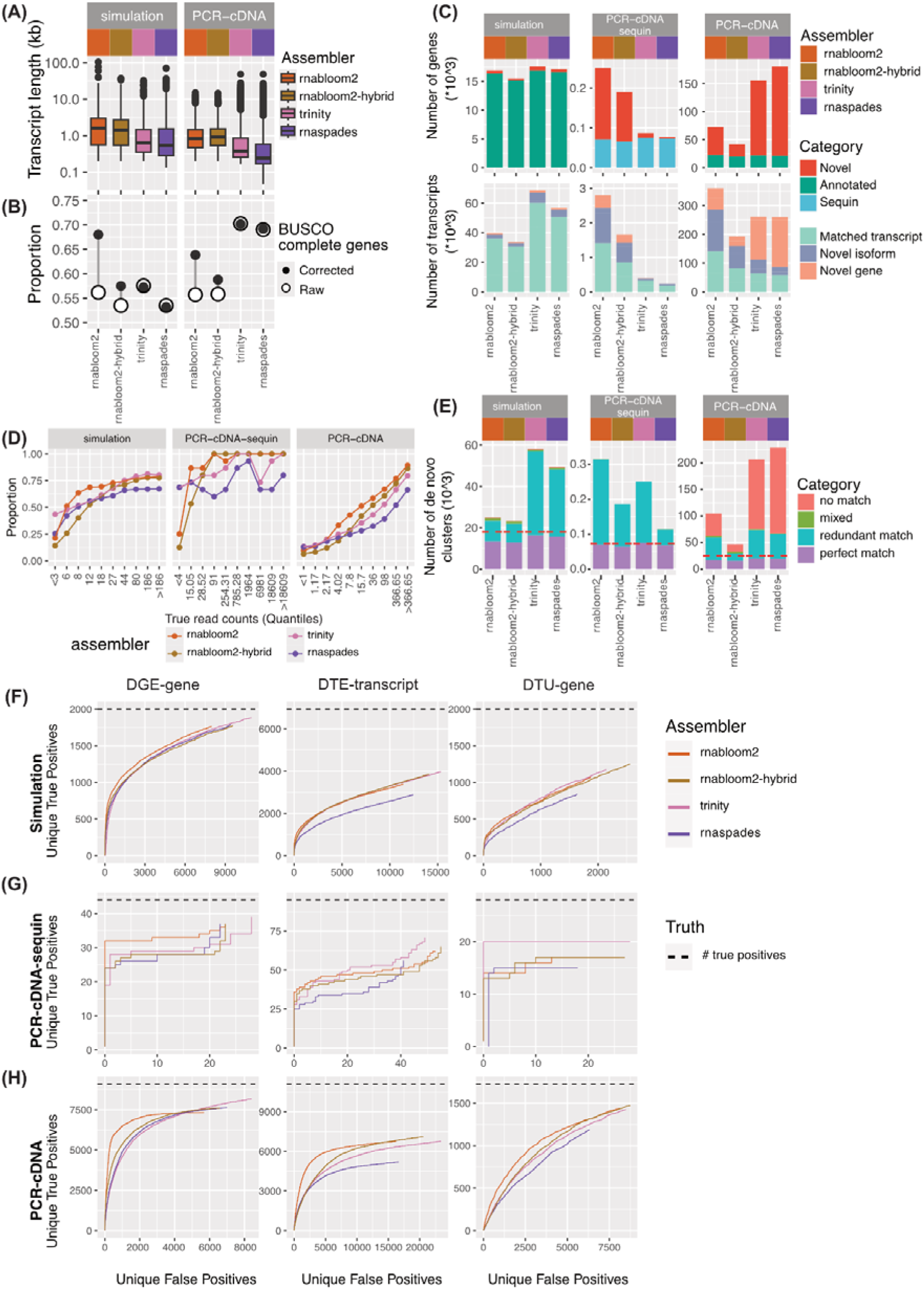
Performance of hybrid short and long-read data assembly. (A) Length distribution of assembled and reference transcriptomes. (B) Proportion of BUSCO complete genes (both single copy and duplicated) assembled before and after sequence polishing. Assembled transcripts were corrected using the reference genome to remove errors and assess protein sequence predictions. (C) Number of novel and annotated genes and transcripts in each assembled transcriptome. Transcripts were further divided into matched transcript (complete splice match and incomplete splice match), novel isoform (novel in catalogue and novel not in catalogue) and novel gene (intergenic, genic, intronic, antisense, and fusion). (D) Proportion of reference transcripts that were assembled binned by true expression. True expression was taken from simulated or Bambu counts, and binned into 10% quantiles. X-axis labels show the corresponding read count range for each quantile. (E) The number of gene-level clusters generated for each assembly. ‘match’ indicates when all transcripts within the cluster correspond to the same reference gene. Where one reference gene matched multiple clusters, the cluster with the most transcripts was classed as a perfect match and the rest as ‘redundant’. Clusters with transcripts from multiple reference genes were classed as ‘mixed’. Clusters with only novel transcripts were classed as ‘no match’. (F-H) Unique false positives were shown against unique true positives for DGE, DTE and DTU analysis for simulation (F), PCR-cDNA-sequin (G) and PCR-cDNA (H) data. Where unique true and positives are described as in Figure 5. The total number of true positives is indicated by the horizontal dashed line (black).

We found that assemblies generated from long reads (RNA-Bloom2) and short reads (Trinity) performed as well as, or better than, the hybrid approaches across most metrics and datasets. RNABloom2-hybrid exhibited similar characteristics to RNA-Bloom2, such as longer transcript lengths (Figure 7A), retained sequencing errors (Figure 7B). However, its overall performance resembled that of RNA-Bloom2 applied to only 30 million reads (Table S3,4,5,6), suggesting that the short-read component of the input data may not have been fully utilised during assembly. Nevertheless, as short reads are used for polishing, RNABloom2-hybrid showed a lower sequence error rate than RNA-Bloom2, as indicated by the smaller improvement observed after error correction when evaluating BUSCO completeness (Figure 7B). RnaSPAdes, by contrast, exhibited features typical of short-read assemblers, producing shorter transcript contigs (Figure 7A) but with low sequence error rates (Figure 7B). We therefore speculate that the long-read component of the input data was underutilised in this case. Collectively, these results suggest that further optimisation is required for hybrid transcriptome assembly approaches to fully exploit the complementary advantages of long- and short-read sequencing technologies.

## Discussion

Our comprehensive evaluation of long-read *de novo* transcriptome assembly tools highlights both the promise and ongoing challenges of performing transcript-level analyses in the absence of a high-quality reference genome and/or transcriptome. We assessed RATTLE, RNA-Bloom2 and isONform, using the short-read assembler Trinity and reference-guided tool Bambu as a guide for expected performance. Using simulated data, sequin spike-ins and real experimental data spanning different library types, read depths, and biological applications, we tested a broad range of experimental scenarios and identified the most effective general strategy for reference-free long-read differential expression analysis.

An important finding of this study is that computational performance is a bottleneck for most tools. Current tools require major improvement to efficiently handle large datasets. This study was one of the first to benchmark these methods with a dataset of a realistic size (60 million reads). However, it is no longer unusual for a PromethION flow cell to generate over 100 million reads, while the recent Pacbio Kinnex SPRQ upgrade on the Revio sequencer now generates 60 million transcript reads per flow cell, and we can only expect the size of long-read transcriptome datasets to increase in the future. Notably, in this study we were unable to assemble the full PacBio single-cell dataset of 555 million reads because of the computational demand. Our strategy of assembling a subset of the data, but leveraging all reads for quantification is one approach for managing the computational burden.

Another key finding is that, although long-read methods inherently produce longer assembled transcripts with more complete isoform structures than short-reads, they still do not reach the capability of reference-guided assemblies. This limitation was seen across multiple metrics, especially in recovering lowly expressed transcripts and performing transcript-level differential analyses.

Among the *de novo* long-read pipelines evaluated, RNA-Bloom2 consistently performed among the top, especially when paired with Corset for transcript clustering and downstream differential analysis. However RNA-Bloom2 had a tendency to produce very large assemblies, with a high amount of redundancy. RATTLE and isONform were both limited by their computational needs, and when applied to a smaller and less noisy dataset, isONform performed well, suggesting its default parameters were not robust on different datasets.

Although Trinity assembled shorter and incomplete transcripts, higher sequencing depth in short-read RNA-seq increased the power in differential testing. Another advantage of short-read RNA-seq is lower error rates in the assembled contigs. We found a surprisingly high rate of sequencing errors were retained in long-read ONT *de novo* assembly. This adversely impacted multiple downstream analysis steps such as clustering and identification of novel transcripts, and we found it to have a profound impact on protein sequence predictions. However, ongoing improvements in basecalling, such as dorado duplex basecalling, and sequencing chemistries, such as the ONT RNA004 kit and R10 flowcell, are expected to reduce these errors. As error rates decrease, we anticipate a corresponding enhancement in transcriptome assembly quality, downstream applications such as DE analysis, as well as an improvement in computational efficiency.

Although comprehensive, our study did not examine ways to improve assembly quality, such as polishing reads or contigs, or combining multiple assemblies produced by different tools or k-mers. Benchmarking could also be expanded to examine alternative methods for post-assembly transcript clustering and quantification, which may influence the final DE results. With the recent development of numerous long-read specific methods, this could improve downstream analysis further (Jousheghani & Patro, 2024; Loving et al., 2024).

We also noted that wider applications of *de novo* transcriptome assembly beyond DE analysis were not benchmarked, but our results have implications for them too. With improved transcript reconstructions, it will become increasingly feasible to perform accurate variant calling and allele-specific expression analyses using long-read data, enabling detailed comparative transcriptomics in non-model organisms. Such analyses can reveal subtle differences across populations or environmental conditions. Although these types of analyses are already performed using short reads, long reads enable phasing of variants and splicing, which can now be done reference-free, and even at single-cell resolution. Additionally, long-read assemblies will allow the study of epitranscriptomic modifications in non-model species. These extensions highlight that assembled transcriptomes are not just a means to perform simple quantification, but can serve as a reference for multiple types of analyses.

Short-read sequencing remains an important approach due to its high throughput, high base quality and well established analytical workflows, allowing the detection of subtle differential expression changes. However, short-read assemblies, as demonstrated, generally produce shorter contigs and are less capable of resolving isoform complexity without a reference. In contrast, long-read assemblies are better at capturing complete transcripts, which is crucial for a comprehensive understanding of gene structure. Thus, the decision to adopt a short-read or long-read strategy, or even a hybrid approach, depends on experimental goals, sample availability, and the quality of the reference genome. Further work is required to identify the optimal and most cost-effective combination of these technologies.

### Conclusion

Our findings demonstrate that while contemporary long-read transcriptome assemblers perform well, weaknesses exist in terms of performance and usage, highlighting the need for ongoing development in this field. By demonstrating the strengths and weaknesses of different combinations of tools, such as RNA-Bloom2 and Corset, we provide a practical guide for researchers designing DE studies without a high-quality reference genome or transcriptome. With future improvements in error-correction methods, assembly algorithms and reference-free quantification strategies, we anticipate that long-read transcriptome assembly will enable more accurate, efficient and wide-ranging applications for long-read data.

## Methods

### Pea growth conditions and RNA extraction

*Pisum sativum* cultivars ‘Thomas Laxton’ and ‘Daisy’ (supplied by the Australian Grains Genebank and the John Innes Centre) were grown within a controlled glasshouse environment at an average ambient temperature of 22°C and subjected to long-day light cycles (8 h dark, 16 h light). The soil medium macro/micro nutrients and other constituents consisted of 0.5 L Vermiculite, 0.5 L Perlite, 35 g Macracote Coloniser Plus 4-month slow release fertiliser (15N : 3P : 9K), 30 g Nitrogen slow-release fertiliser (40N : 0P : 0K), 25 g Water holding granules, 15 g Trace elements (6Mg : 6.5FE :5.4S : 1.5Mn : 0.4Zn : 0.14B : 0.07Mo), and 5 g Garden lime per 30 L of soil. Whole stipules were harvested from the third leaflet node of individual seedlings 2 weeks post-emergence and snap-frozen in liquid nitrogen. Each sample was crushed into a fine powder by pestle and mortar. Samples were then weighed on a microgram scale to 1 mg of whole stipule tissue per extraction. Total RNA was then isolated using Invitrogen TRIzol reagent (ThermoFisher Scientific) using the manufacturer’s protocol. Minor modifications to the protocol included an extended centrifugation for 15 minutes, as some RNA precipitate was still suspended in solution for some samples. RNA precipitate pellets were washed twice rather than once, as it was found this removed further impurities and improved the overall quality of the RNA. Centrifugation between washes was also extended to 10 minutes, to ensure the RNA pellet was anchored to the bottom of the Eppendorf tube during wash steps. Total RNA pellets for each sample were then re-suspended in RNase-free water, and subsequently quality-checked and quantified utilising the NanoDropTM spectrophotometer.

### Simulating Nanopore cDNA data and matched short-read sequencing data

We first obtained a subset of transcripts that are widely expressed in the GTEx v9 long read dataset (92 samples) using Gencode comprehensive annotation (v44). We kept transcripts with more than 5 reads in at least 15 samples after Salmon quantification (18145 genes, 40509 transcripts) and stored their mean count per million (CPM) values as the control group’s baseline expression. We then generated a perturbed set of CPM values where transcript expression was changed by: (1) randomly selecting 1000 genes and changing all transcripts belonging to that gene concordantly (500 genes 2 fold up and 500 genes 2 fold down), (2) selected another 1000 genes randomly, and then select 2 random transcripts from the gene and swap their expression, (3) selected another 1000 genes randomly and then selected 1 random transcript to change its expression (500 transcripts 2 fold up and 500 transcripts 2 fold down). The updated CPM were stored as the perturbed group baseline expression. We then generated a count matrix and CPM matrix for 3 control replicates and 3 perturbed replicates with gamma distribution, followed by a Poisson distribution (Baldoni et al., 2024). Both long-read and short-read FASTQ files were simulated using SQANTI-SIM with default settings and ONT R9.4 cDNA error profile (v 0.2.1) (Mestre-Tomás et al., 2023). The long read data contained 6 million reads in total, and an average read length of 1085 bp, and short read data was 100 bp paired-end. We then subsampled the short-read data to match the total number of base pairs in the long read data (6.5 billion bases). The simulated data was non-stranded, and contains 2000 DE genes, 2000 genes with DTU, 5927 transcripts with DTU and 6933 DE transcripts. It is available for download at https://doi.org/10.5281/zenodo.14263456.

### Nanopore PCR-cDNA, direct-RNA data and matched short-read RNA-seq

To investigate how the assemblers perform with different sizes and types of data, we downsampled the PCR-cDNA data for 2 cancer cell lines (HCC827 and NCI-H1975, 3 replicates each) from GSE172421 to 12 million, 30 million and 60 million reads in total (2 million, 5 million and 10 million for each replicate, with a mean read length 685 bp) and included short-read stranded RNA-seq with matched number of bases to the 60 million PCR-cDNA data (75bp paired-end and 41 billion bases) (Dong et al., 2023). The PCR-cDNA data includes sequin spike-in transcripts from 2 mixes of different concentrations. Our benchmarking study also used direct-RNA data from the SG-NEx project for the A549 and MCF7 cell lines (3 replicates each, 11.6 million total long reads with a mean read length of 1060 bp, covering 12.3 billion bases) and matched short-read RNA-seq (Chen et al., 2021). Lastly, a PCR-cDNA ONT dataset with 12 samples (3 replicates across 4 conditions) from pea (*Pisum sativum*) species was prepared with the Oxford Nanopore PCR-cDNA Barcoding kit (SQK-PCB109) following the manufacturer’s guidelines. Fifty-nanograms total RNA input was used for each of the 12 samples. Samples were reverse-transcribed into cDNA and PCR amplified and barcoded for 15 cycles in a thermocycler. The final pool was loaded onto a PromethION flow cell (FLO-PRO002, R9.4.1) and sequenced on the PromethION P24 (Oxford Nanopore Technologies Limited) over 72 hours. Raw nanopore reads were basecalled in super-accurate mode and demultiplexed with guppy (v 6.3.7). This data contained 16.6 million reads covering 10.7 billion bases. The same RNA was used to prepare short-read libraries with the Illumina TruSeq stranded kit v2 according to manufacturer’s instructions. Pooled libraries were sequenced on an Illumina NextSeq 550 with 75 bp single end reads. All long-read PCR-cDNA data, dRNA data and their matched short read were stranded.

### PacBio Kinnex PBMC single cell RNA-seq data

PacBio Kinnex PMBC single cell RNA-seq data was downloaded from https://downloads.pacbcloud.com/public/dataset/Kinnex-single-cell-RNA/. The deconcatenated and UMI-deduplicated FASTQ files for 3 10x 3’ PBMC samples were downloaded (10k, 20k_rep1, 20k_rep2). The full data were subsampled to 5 million reads (mean read length 835bp) per sample for assembling. The full dataset containing 555 millions reads was used for quantification and differential analysis.

### Transcriptome assembly

RATTLE (v 1.0), RNA-Bloom2 (v 2.0.1), isONform (v 0.3.4) and Trinity (v 2.15.1) were run using FASTQ files according to the default workflow. RATTLE was compiled from source, RNA-Bloom2 and isONform were installed from conda, and Trinity was run within a singularity container image. Simulated cDNA data were assembled using an unstranded setting, while the PCR-cDNA data from cancer cell lines and pea were preprocessed with pychopper (v 2.7.9) to reorientate reads retaining only full length and rescued reads with properly paired primers at both ends, and assembled using a stranded setting. Direct RNA was assembled with a stranded setting, and U was converted to T in the final assembly. The isONform pipeline was run with isONclust (0.0.6.1, with the `--ont` option), isONcorrect (0.1.3.5, default options) and isONform (`--exact_instance_limit 50 --k 20 --w 31 –xmin 14 --xmax 80 --max_seqs_to_spoa 200 --delta_len 10 --iso_abundance 5 --delta_iso_len_3 30 --delta_iso_len_5 50`). For the single-cell PacBio data, we ran RATTLE in stranded mode, RNA-Bloom2 in stranded mode with `–lrpb` flag, and isONform pipeline without the correction step suggested by the authors. Details can be found in the GitHub repository https://github.com/DavidsonGroup/longread_denovo_benchmark.

For reference-guided assembly, minimap2 (v 2.26-r1175) was used to generate BAM files (`-ax splice` for simulated cDNA, `-ax splice -uf` for PCR-cDNA data from cancer cell lines and pea and `-ax splice -uf -k14` for dRNA). For simulation, cell lines and single-cell data the GRCh38 genome and Gencode v44 comprehensive GTF annotation were used (with sequin annotation included for PCR-cDNA data). For the pea data, the RefSeq ZW6 reference genome and corresponding gene annotation were used (Table S1).

Bambu (v 3.3.4) was then run with the corresponding strandness of the data and other parameters default. Only expressed transcripts and genes (based on a Bambu transcript count of >= 1 in at least one sample) were retained for downstream comparisons and differential analysis. The Bambu GTFs were converted to FASTA using gffread (v 0.12.8).

All assemblies were executed on a high-performance cluster at the Walter and Eliza Hall Institute, Australia, with 48 cores (Intel(R) Xeon(R) CPU E5-2690 v4 @ 2.60GHz (Broadwell), Intel(R) Xeon(R) Gold 6342 CPU @ 2.80GHz (Icelake), Intel(R) Xeon(R) Gold 6130 CPU @ 2.10GHz (Skylake)) and 1Tb max RAM (DDR4 2400 MT/s, DDR4 3200 MT/s, DDR4 2666 MT/s) requested. Assemblies that did not finish in 2 weeks were terminated and excluded.

### Hybrid assembly

To perform hybrid assembly on the simulation data and 60 million PCR-cDNA data, we subset 50% of Nanopore reads and 50% of Illumina reads. RNA-Bloom2 (hybrid mode) and rnaSPAdes (4.2.0, binary downloaded from github) were used for hybrid assembly with the same strandness for each dataset as described earlier.

### Assessing basic assembly quality metrics

We assessed all assemblies with transrate (v 1.0.3) for assembly length, GC content and reference recovery using Conditional Reciprocal Best BLAST (Smith-Unna et al., 2016). We also run BUSCO (v 5.4.7, database `primates_odb10` and `fabales_odb10`) to obtain the proportion of conserved genes that were assembled (Manni et al., 2021). We used RepeatMasker (4.2.2) to search repetitive sequences in human and Pisum sativum databases (Dfam 3.9) (Storer et al., 2021). We used PrimeSpotter to detect internally primed sequences from the assembled transcriptome (You et al., 2025).

### Assessing assembly based on splice junction using SQANTI3

We ran SQANTI3 (v 5.1.2) to assign *de novo* assembled transcripts and Bambu novel transcripts to reference gene and transcript annotation (Pardo-Palacios, Arzalluz-Luque, et al., 2024). We provided a junction file to correctly align assembled transcripts to the reference genome. The aligned BAM files were converted to GTF and then FASTA files based on reference genome sequences using gffread to remove small insertion and deletions. These genome-corrected FASTA files were also assessed for their BUSCO score. The original uncorrected FASTA files were used for all downstream analysis. For simulated non-stranded data, all transcripts classified as ‘antisense’ were subjected to a second round of SQANTI3 annotation using their reverse complement sequences.

### Assessing the recovery of reference transcripts and genes

To measure the recovery of reference transcripts and genes, we binned the reference transcripts and genes into 10 quantiles based on quantification from simulation, sequin or Bambu with reference. The proportion of assembled reference transcripts and genes was extracted from SQANTI3 results.

### Assessing transcript quantification correlation

To quantify the expression of assembled transcripts, we ran minimap2 (map-ont for ONT data and map-hifi for PacBio data) to align the FASTQ files from individual samples to the *de novo* transcriptome (options: `-p 1.0 -N 100`). For the ONT and short-read data, we then ran salmon (v 1.10.2) on the aligned BAMs with 100 bootstraps, for the ONT data we used the option `--ont`. For the hybrid assembly data, the counts from short read (salmon quant) and long reads (minimap2+salmon) were summed to obtain the final counts for each assembled transcript in each replicate. For the single-cell data, oarfish were used and single-cell counts were aggregated to sample level before assessing quantification. To measure correlation with truth transcript expression, *de novo* transcripts were assigned to reference transcripts based on SQANTI3 results. When multiple *de novo* transcripts were assigned to the same reference transcript, only the one with the highest expression was used to calculate the correlation. *De novo* transcript expression was calculated as log2(count +1). The expression values for each sample were then correlated to known expressions for simulation (log2(count +1)), sequin data (log2(concentration)), or Bambu quantification (log2(count +1)).

### Assessing transcript to gene cluster assignment

Because Bambu, isONform, RATTLE, Trinity and rnaSPAdes perform clustering of transcripts to genes using their in-built algorithms, we included their transcript to gene cluster information. For all *de novo* methods, we further clustered assembled transcripts into genes using Corset based on the all-to-all overlap of transcripts using minimap2 (with options: `-a -k15 -w5 -e0 -m100 -r2k -P --dual=yes --no-long-join`) (Davidson & Oshlack, 2014). Furthermore, the performance of clustering was evaluated using several metrics including precision (*de novo* transcripts associated with transcripts from the same gene are clustered into one gene cluster) and recall (*de novo* transcripts associated with transcripts from the different genes are separated into different gene clusters) using SQANTI3 assigned transcripts and genes as truth.

### Assessing gene cluster quantification

We use 3 different methods to obtain gene clusters for quantification: (1) *de novo* gene clustered from Corset, (2) *de novo* gene clusters from native clustering if applicable, (3) gene clusters defined by SQANTI3 using the reference gene annotation. Gene cluster expression was calculated as aggregated counts from all transcripts in the cluster and converted to log2(count+1). Using SQANTI3 results, each gene cluster was assigned to a known gene if the majority of its transcripts were assigned to that gene. The correlation was then evaluated in a manner similar to that used for transcripts.

### Assessing the redundancy of gene clusters

For *de novo* gene clusters from Corset or native clustering, we further classified them into (1) no match: if all transcripts from the cluster have no match to reference; (2) mixed: if transcripts from the cluster match to multiple reference genes, (3) match: all transcripts from the cluster match to one reference gene, and if multiple clusters match to the same gene, the one with most transcripts are defined as a perfect match and the rest defined as a redundant match.

### Assessing differential analysis using *de novo* assembly

For ONT and short read assemblies, we used edgeR::catchSalmon (4.2.0) to import counts and read to transcript ambiguity. The scaled transcript counts (counts divided by read-to-transcript ambiguity) were used for differential transcript analysis (DTE and DTU), and the sum of transcript counts from gene clusters was used for differential gene expression analysis (DGE). For hybrid assemblies, the scaled transcript counts from short and long read were aggregated for DTE/DTU analysis and the sum of short and long read transcript counts from gene clusters was used for differential gene expression analysis (DGE). For single-cell data, due to the sparsity of the data, we performed pseudo-bulk differential analysis at cluster level. First, we obtained true cell type labels. We downloaded the Seurat count matrix for the 3 samples from the PacBio website and annotated with Azimuth (0.5.0) using PBMC reference data to obtain true cell types for each cell (Hao et al., 2021). Then for each of the *de novo* assemblies, a single cell UMI count matrix was generated using oarfish (Zare Jousheghani et al., 2025), and transcript UMI counts were aggregated into gene UMI counts based on gene clusters. Gene expression clustering was then performed to confirm the presence of two major clusters, corresponding to lymphoid and monocytic cells based on true cell type labels, and pseudo-bulking was performed for the two major clusters in each replicate. All analyses were performed using the limma::voom (3.52.4) pipeline after filtering (Baldoni et al., 2024). For the single-cell data, we also ran limma::treat to select differential genes and transcript with fold-change >1.5. For differential transcript usage (DTU) analysis, we used limma::diffSplice, and Simes correction to obtain gene-level p values. The differential lists were compared to the truth of either simulated differential gene/transcript, sequin with known differential expression or abundance, or Bambu differential results with the reference. If multiple *de novo* transcripts or genes matched to a reference transcript or gene, only the one with the smallest FDR was used when assessing unique true or false positives. The code for differential analysis can be found in the GitHub repository https://github.com/DavidsonGroup/longread_denovo_benchmark.

### Novel transcripts in the *de novo* assembly

We ran JAFFAL (v 2.3) on the RNABloom2 and RATTLE assembled transcriptome in FASTA format from the cancer cell lines (PCR-cDNA and dRNA) (Davidson et al., 2022). Known structural variants and fusions from corresponding cell lines were downloaded from the DepMap portal (Public 24Q2 and CCLE2019 dataset) (Ghandi et al., 2019; Tsherniak et al., 2017). Alignment and tracks were visualised in IGV (Robinson et al., 2011).

## Data access

The code to reproduce the analysis can be found in GitHub: https://github.com/DavidsonGroup/longread_denovo_benchmark. The simulated data can be downloaded from https://doi.org/10.5281/zenodo.14263456. The PCR-cDNA data from cancer cell lines were obtained from GEO (GSE172421), and dRNA data were obtained from SG-NEx AWS Open Data Registry: https://registry.opendata.aws/sgnex/. The long-read PCR-cDNA and short-read RNA-seq data from pea are available in ArrayExpress under accessions E-MTAB-14841 and E-MTAB-14845 respectively. The PacBio Kinnex single cell RNA-seq data were downloaded from https://downloads.pacbcloud.com/public/dataset/Kinnex-single-cell-RNA/. All assemblies generated in this study can be obtained from https://doi.org/10.5281/zenodo.17538009

## Acknowledgement

This research was undertaken with the assistance of Milton HPC/Virtual Machines, supported by WEHI Research Computing. We thank Ashley L. Weir for her valuable comments on the manuscript. We also acknowledge the use of ChatGPT for assistance with language and grammar editing.

## Author contribution

FY and NMD developed the conceptual design of the benchmarking project. FY performed all analyses and generated figures. FY and NMD wrote and revised the manuscript. PLB designed the simulation dataset with differential expression. JL and QG collected *Pisum sativum* samples, prepared libraries, performed sequencing, provided valuable suggestions for the benchmark and manuscript. NMD, MGL and MER supervised data generation and analyses. All authors provided feedback and approved the final manuscript.

