## Supplementary Figures (S1-5) for "Towards accurate, reference-free differential expression: A comprehensive evaluation of long-read *de novo* transcriptome assembly"

Table S1, Summary of dataset characteristics, parameters and analyses done for each dataset.

Table S2, Running time and memory usage for each assembler in each dataset.

Table S3, The number of transcripts and genes by SQANTI3 category for each assembly.

Table S4, Number of transcripts matching BUSCO genes for each assembly using (1) the raw uncorrected transcriptome sequences and (2) genome-corrected transcriptome sequences.

Table S5, Percentage of repetitive sequences and internally primed sequences.

Table S6, Correlation coefficient between *de novo* assembly and reference transcripts and gene abundances.

Table S7, Clustering evaluation metrics of native and Corset clustering using SQANTI3 to define true clusters.

Table S8, The number of clusters classes as perfect, redundant, mixed and no match for each clustering method.

Table S9, Top 10 novel DTEs ranked by FDR in 60 million PCR-cDNA data.

Table S10, Top 10 novel DTEs ranked by FDR in dRNA data.

Table S11, Novel fusion DTEs in 60 million PCR-cDNA data.

Table S12, Novel fusion DTEs in dRNA data.

Table S13, Novel DTE with BUSCO hits in pea data.

### Supplementary figures S1-S5 for:

Towards accurate, reference-free differential expression: A comprehensive evaluation of long-read *de novo* transcriptome assembly

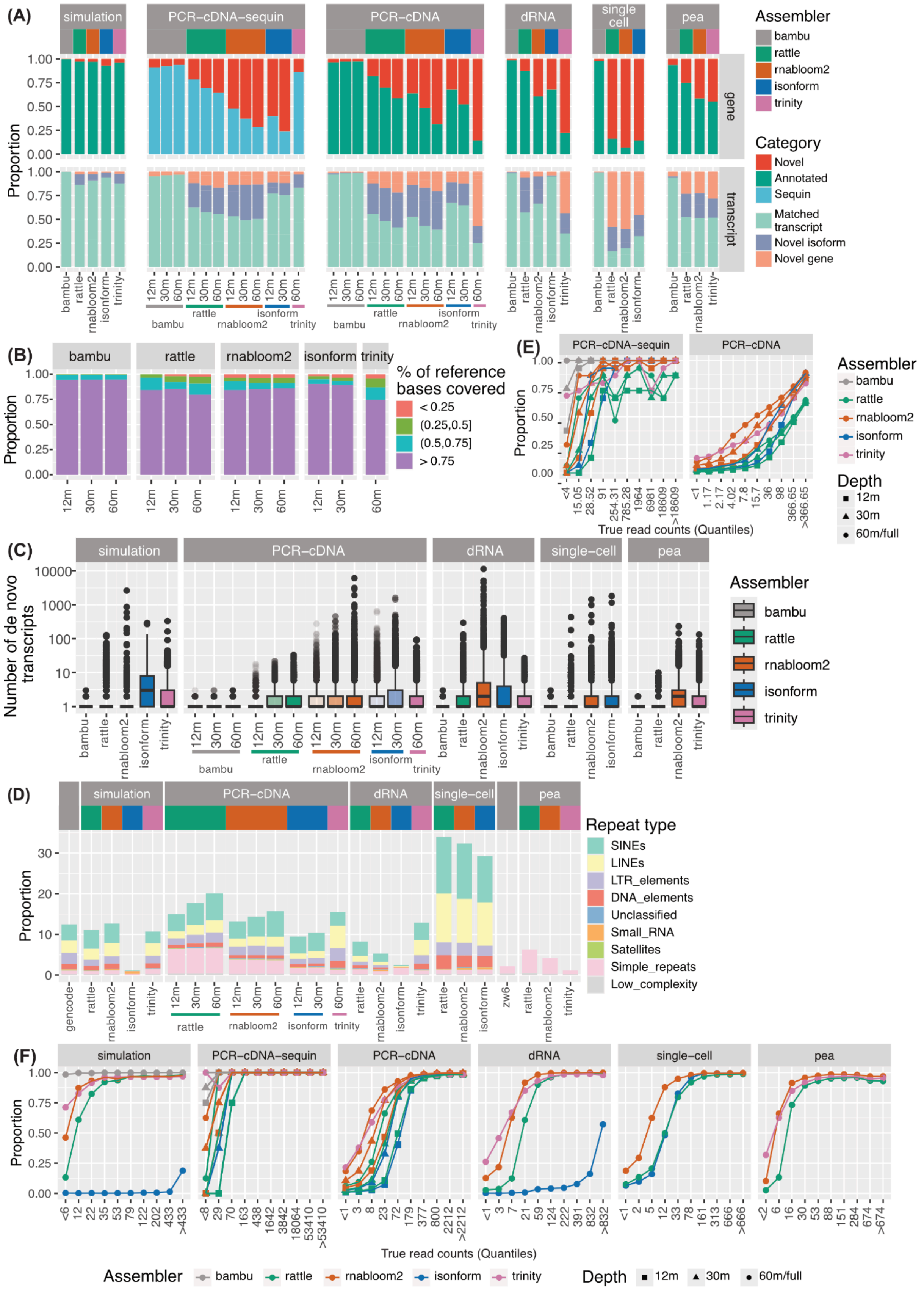


*Figure S1: Assessment of assembled transcriptome quality*

*(A) Proportion of novel and annotated genes and transcripts in each assembled transcriptome. Transcripts were further divided into matched transcript (complete splice match and incomplete splice match), novel isoform (novel in catalogue and novel not in catalogue) and novel gene (intergenic, genic, intronic, antisense and fusion). Results were generated by running SQANTI on each assembly.*

*(B) The proportion of bases recovered for each sequin transcript in PCR-cDNA data using the Conditional Reciprocal Best BLAST (CRBB) approach.*

*(C) Number of de novo transcripts per reference transcript*

*(D) Proportion of repetitive elements colored by repeat types in each assembly and reference (grey boxes). Gencode denotes the human gencode v44 annotation. Zw6 denotes the pea GCF_024323335.1_CAAS_Psat_ZW6_1.0 annotation.*

*(E) Proportion of reference transcripts that were assembled, binned by true expression. True expression was taken from simulated or Bambu counts and binned into 10% quantiles. X-axis labels show the corresponding read count range for each quantile. Read depth is indicated by the symbol shape for PCR-cDNA.*

*(F) Proportion of reference genes that were assembled, binned by true expression. True expression was taken from simulated or Bambu counts, and binned into 10% quantiles. X-axis labels show the corresponding read count range for each quantile. Read depth is indicated by the symbol shape for PCR-cDNA.*

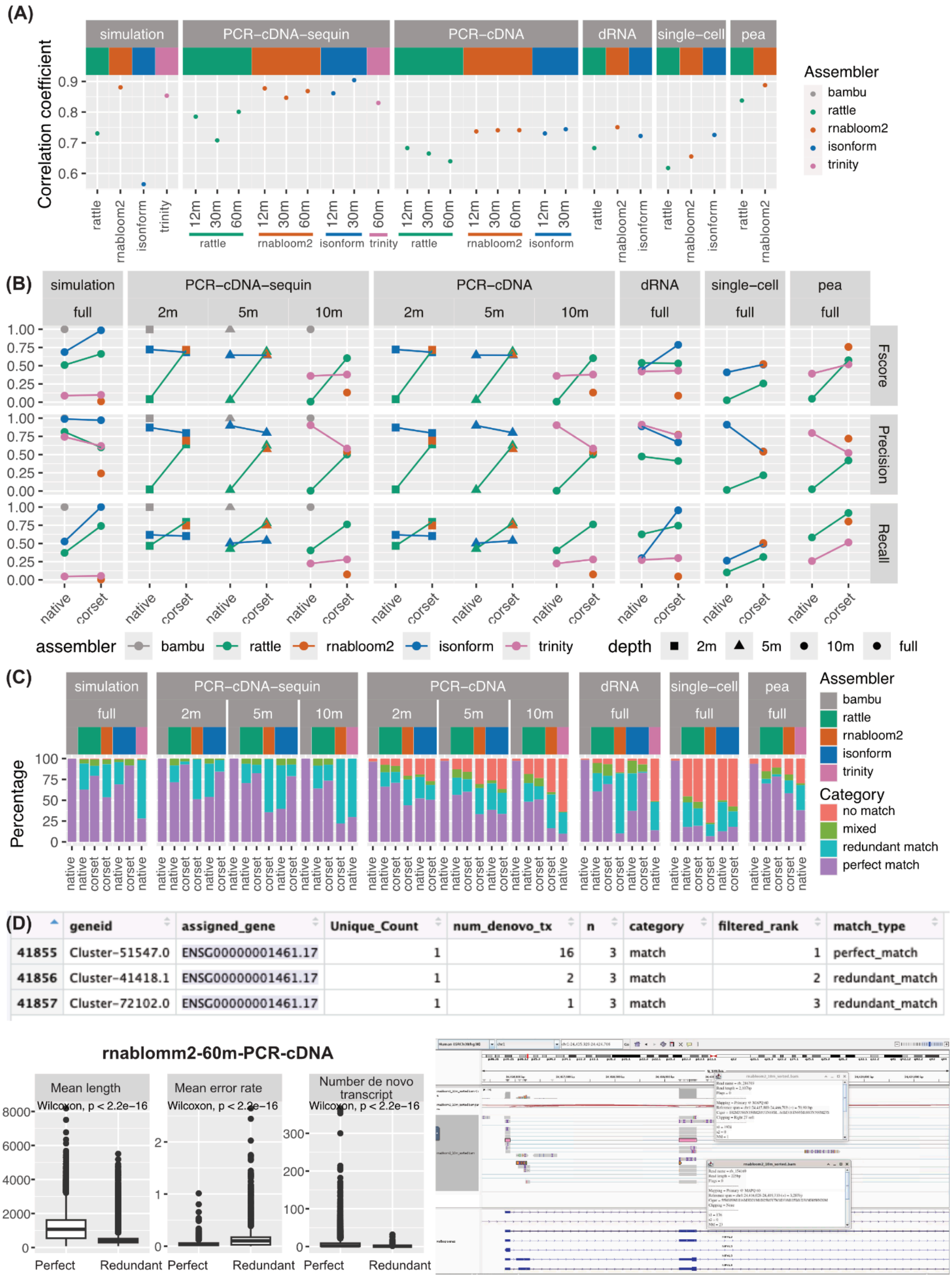


*Figure S2: Accuracy of transcript and gene abundance estimates.*

*(A) Pearson correlation coefficients between estimated and true transcripts expression values (log2(count+1)) excluding unassembled reference transcripts.*
*(B) The precision, recall and F1 score for native and Corset clusters. True clusters were defined using SQANTI3 assigned genes. We note that clustering metrics are not directly comparable between assemblies due to their differences in the number of transcripts and gene clusters. Assemblies that capture more complexity may suffer from lower scores.*
*(C) The proportion of gene-level clusters generated for each assembly. 'match' indicates when all transcripts within the cluster correspond to the same reference gene. Where one reference gene matched multiple clusters, the cluster with the most transcripts was classed as a perfect match and the rest as 'redundant'. Clusters with transcripts from multiple reference genes were classed as 'mixed'. Clusters with only novel transcripts were classed as 'no match'.*
*(D) Top: Example of an RNA-Bloom2 redundant cluster corresponding to the same reference gene (ENSG00000001461.17) from PCR-cDNA 60 million data. Each row is a de novo gene cluster from Corset. Geneid is the Corset gene cluster ID. Assigned_gene is the SQANTI3 assigned gene. Unique_count is the number of unique assigned genes in each cluster. Num_denovo_tx is the number of de novo transcripts in each cluster. N represents the number of occurrences of the same assigned gene.*
*Bottom left: Box plots showed mean transcript length, mean error rate and number of transcripts comparing perfect cluster to redundant cluster in RNA-Bloom2 assembly from the PCR-cDNA 60 million data.*
*Bottom right: IGV visualization showing a transcript from the redundant match cluster, Cluster-72102.0 (orange) has a higher error rate (23 mismatches vs 1 mismatch) and is shorter (225 bp vs 2107 bp) compared to a transcript from the perfect match cluster, Cluster 51547.0 (pink).*

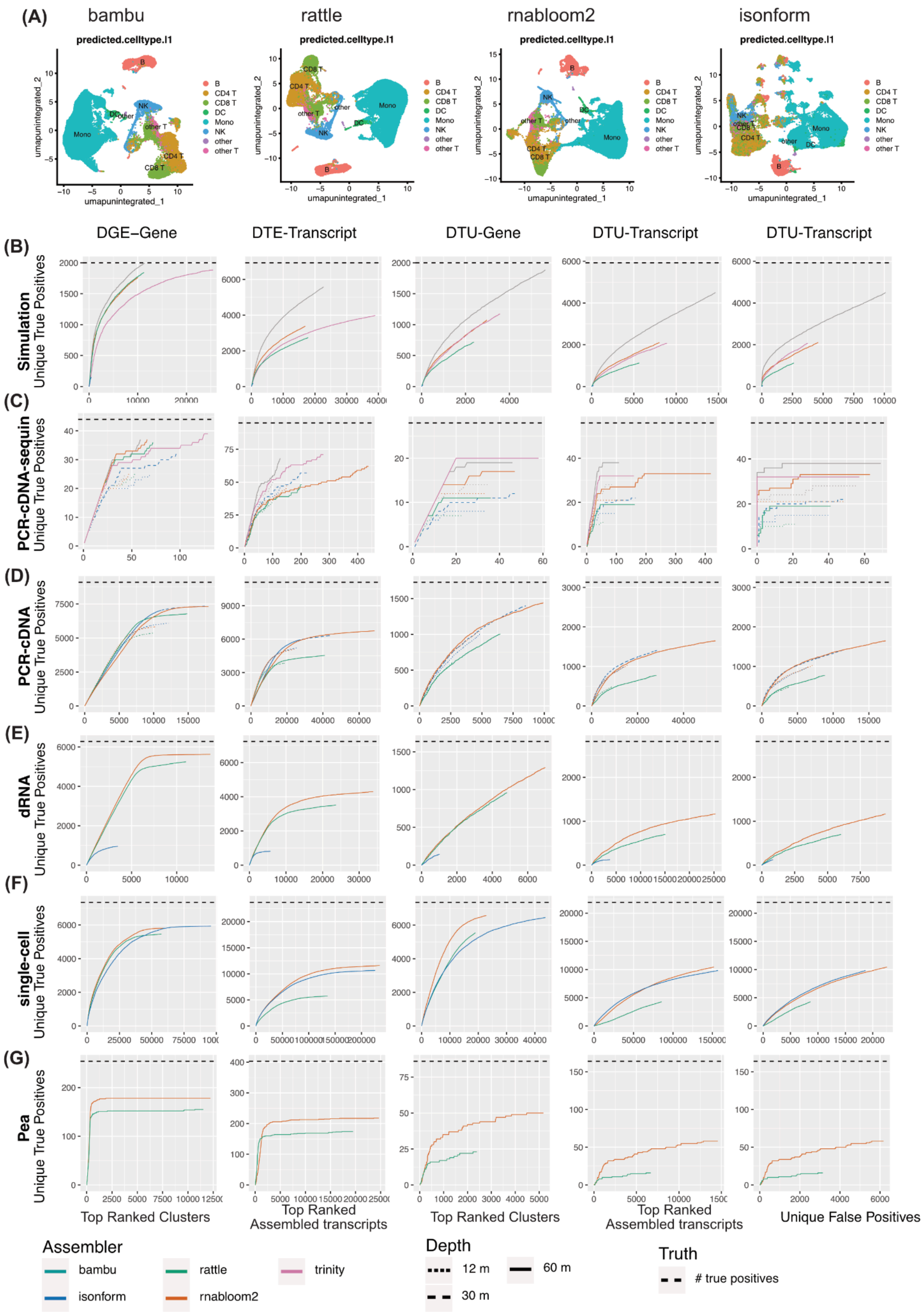


*Figure S3 ROC-style curves for differential analysis.*

*(A) UMAP generated using transcript-level counts from reference (bambu) and de novo assembly and colored based on the cell type annotation based on count matrix from PacBio.*

*(B-G) DGE, DTE and DTU analysis were performed for the (B) Simulation, (C) PCR-cDNA sequin, (D) PCR-cDNA, (E) dRNA, (F) single cell and (G) pea dataset. The number of accumulative true positives is shown as a function of rank after ordering by FDR (columns 1-4) or as a function of accumulative false positives (column 5). When multiple clusters match the same reference gene, the cluster with the lowest FDR is retained, hence 'unique' true positives and 'unique' false positives. Truth is defined from simulation and sequin or Bambu differential results for the rest, and the total number of true positives is indicated by the horizontal dashed line. For panels (C-D), the dotted and solid lines show the number of true positives and false positives from the 12 million and 60 million PCR-cDNA datasets, respectively.*

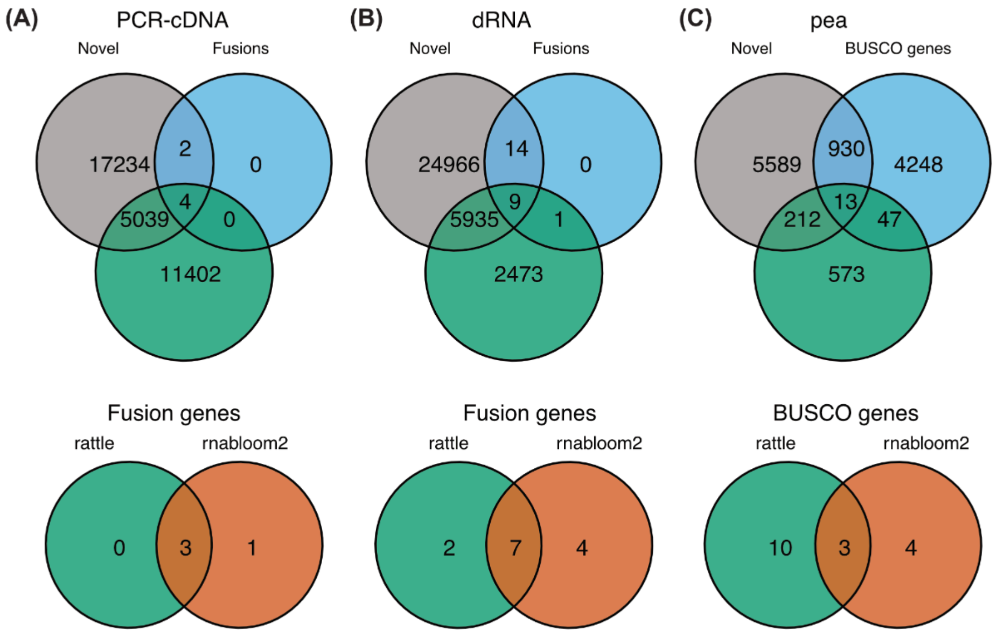


***Figure S4 Novel transcript in RATTLE***

*(A) Top: Venn diagram showing the number of transcripts from RATTLE which were: differentially expressed, novel and/or known fusion genes in PCR-cDNA 60 million data. Bottom: overlap of CCLE fusion genes assembled by RNA-Bloom2 and RATTLE in PCR-cDNA 60 million data.*

*(B) Top: Venn diagram showing the number of transcripts from RATTLE which were: differentially expressed, novel and/or known fusion genes in dRNA data. Bottom: overlap of CCLE fusion genes assembled by RNA-Bloom2 and RATTLE in dRNA data.*

*(C) Top: Venn diagram showing the number of transcripts from RATTLE which were differentially expressed, novel, and/or had a BUSCO gene match in pea data. Bottom: overlap of BUSCO genes assembled by RNA-Bloom2 and RATTLE in pea data.*

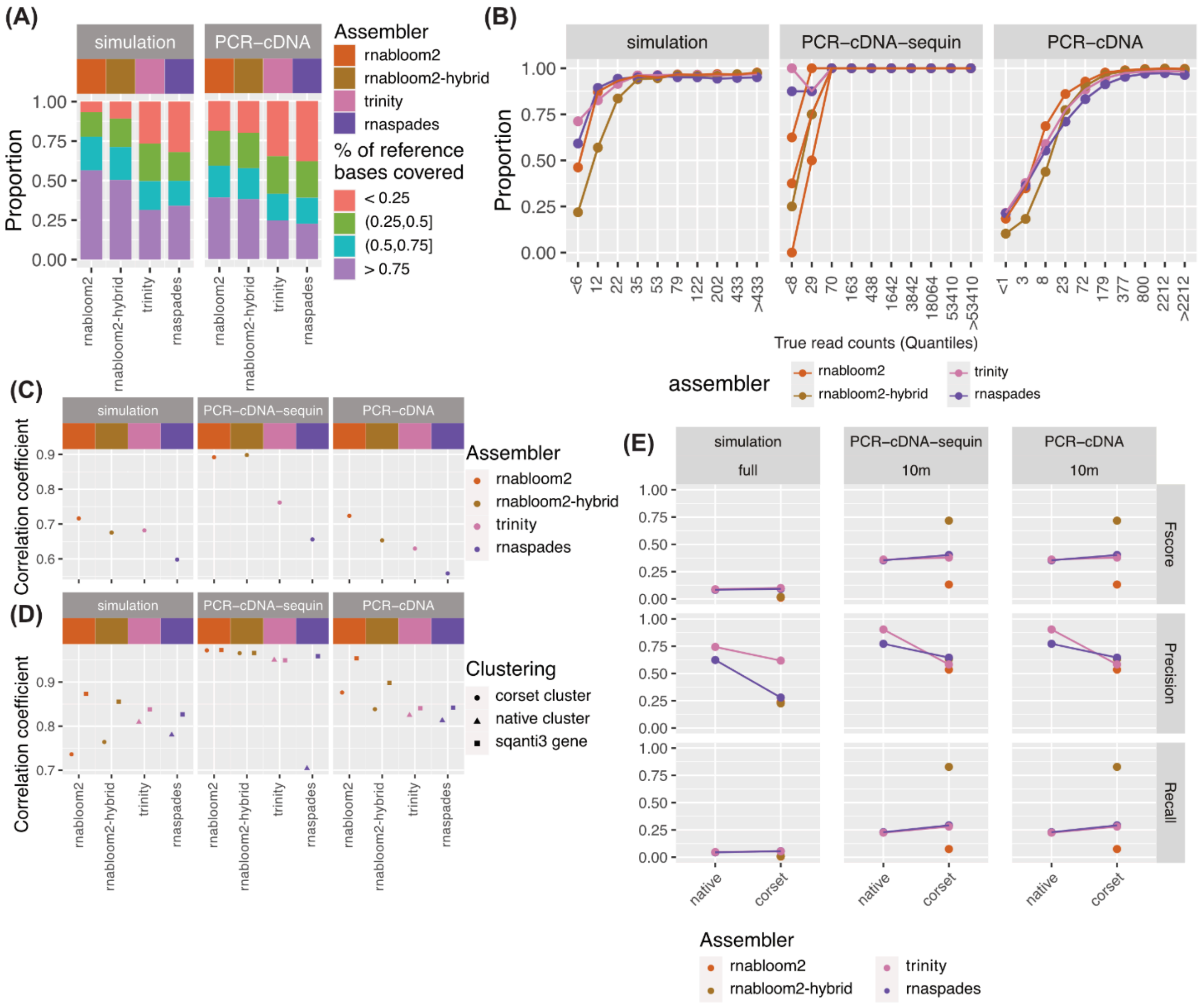


***Fig S5 Hybrid de novo assembly***

*(A) The proportion of bases recovered for each reference transcript using the Conditional Reciprocal Best BLAST (CRBB) approach.*

*(B) Proportion of reference genes that were assembled, binned by true expression. True expression was taken from simulated or Bambu counts, and binned into 10% quantiles. X-axis labels show the corresponding read count range for each quantile. Read depth is indicated by the symbol shape for PCR-cDNA.*

*(C) Pearson correlation coefficients between estimated and true transcripts expression values (log2(count+1)).*

*(D) Pearson correlation of true gene expression (log2(count+1)) to Corset, native or SQANTI3 cluster expression. SQANTI3 clustering is an optimal scenario where transcripts are grouped based on their true reference genes, and it is included as an upper limit.*

*(D) The precision, recall and F1 score for native and Corset clusters. True clusters were defined using SQANTI3 assigned genes.*
